# EGP1K: Whole-Genome Sequencing of 1,024 Egyptians Characterizes Population Structure and Genetic Diversity

**DOI:** 10.64898/2026.04.02.715521

**Authors:** Khaled Amer, Ahmed Moustafa, Wael A. Hassan, Eman Adel, Khlood R. AbdElaal, Tasnim A. Ghanim, Ahmed Abd El-Raouf, Ahmed El-Hosseiny, Ahmed F. El-Sayed, Ahmed H. Badr, Alaa Hassan, Amira Kotb, Amira Ragheb, Ashraf M. Muhammad, Asmaa Ali, Ayten Abdelaal, Eman Ramadan, Fadya M. El-Garhy, Hasnaa El Shehaby, Mahmoud A. Ali, Mahynour Albarbary, Marwa A. Zahra, May Amer, Mohamed A. Elmonem, Nermeen T. Fahmy, Omnia M. Abdel-Haseeb, Tokka M. Hassan, Yara A. Daoud, Yasmeen Howeedy, Yasmeen K. Farouk, Sameh Soror, Gina El-Feky, Mahmoud Sakr, Neveen A. Soliman, Yehia Z. Gad, Khaled A. Abdel-Ghaffar, Egypt Genome Consortium

## Abstract

Middle Eastern and North African populations remain underrepresented in genomic databases, comprising less than 1% of genome-wide association study participants despite representing approximately 6% of the global population. Here we present the Egypt Genome Project (EGP1K), in which we performed whole-genome sequencing on 1,024 unrelated Egyptian individuals originating from 21 of Egypt’s 27 governorates, recruited through eight clinical and research centers across Upper and Lower Egypt.

We identified over 51.3 million variants, of which 17.1 million (33.4%) were absent from dbSNP. Allele frequency comparisons across 6.5 million shared variants showed the strongest concordance with Middle Eastern populations (*τ* = 0.977). Principal component analysis and ADMIXTURE modeling at K = 7 revealed that Egyptians share a dominant ancestry component (71.8%) with Middle Eastern populations and carry a smaller Egyptian-enriched component (18.5%) that distinguishes them from neighboring groups. Runs of homozygosity varied substantially across subregions, with Upper Egypt showing the highest burden, paralleling elevated consanguinity rates. Carrier frequency analysis identified *MEFV* (Familial Mediterranean Fever) at 9.1% as the most prevalent pathogenic carrier state; when adjusted for the national consanguinity rate, *MEFV* carrier status alone projects approximately 6,600 affected births per year. HLA class I typing identified allele frequencies placing Egyptians within the Levantine-Eastern Mediterranean cluster, providing baseline immunogenetic data currently absent from international databases.

Analysis of polygenic risk score distributions revealed substantial differences in threshold-based risk stratification between Egyptians and European reference populations. When the European-derived 90th percentile threshold was applied, 83.3% of Egyptians were assigned to high-risk strata for stroke, 76.4% for chronic kidney disease, and 72.8% for gout, compared to the intended 10% high-risk proportion. These distributional shifts were observed across several cardiometabolic traits (Cohen’s d = 1.55-1.61), while other traits showed closer cross-population concordance, indicating that the degree of threshold miscalibration varies by trait.

Together, these findings establish EGP1K as a genomic reference for Egypt and indicate that European-derived risk stratification thresholds may not be directly transferable to the Egyptian population, supporting the need for population-specific calibration of polygenic risk scores.

## Introduction

Population genomic studies have identified variants associated with disease susceptibility, drug response, and evolutionary adaptation across diverse human groups. However, the clinical utility of these findings depends on adequate representation of study populations in reference databases. The 1000 Genomes Project, the Human Genome Diversity Project, and subsequent initiatives have sequenced thousands of human genomes, yet Middle Eastern and North African populations remain among the most underrepresented groups (Bhattacharya et al. 2023). Middle Eastern populations comprise less than 1% of participants in genome-wide association studies, a disparity that limits the transferability of genomic findings to these populations (Martin et al. 2019; Need and Goldstein 2009; Popejoy and Fullerton 2016; Sirugo, Williams, and Tishkoff 2019).

Egypt represents one of the most significant gaps in genomic representation within the MENA region. With over 100 million inhabitants, Egypt is the most populous nation in the Middle East and North Africa (MENA) region, yet its genomic architecture has been characterized only through limited regional sampling. Wohlers et al. (2020) reported the first Egyptian genome reference using 110 individuals, identifying population-specific insertions and haplotypic expression patterns. The present study builds on this foundation with a larger, geographically distributed cohort.

Egypt lies at the junction of Africa, Asia, and Europe, a location that has shaped its genetic composition over millennia of human migration and cultural exchange. This geographic position suggests a complex population structure that cannot be captured through geographically restricted sampling. Egypt also displays marked regional variation in consanguinity practices. Populationbased surveys estimate overall consanguinity rates of 29 to 36%, with substantially higher prevalence in rural areas (up to 60%) than in urban centers (approximately 17%) (Hafez et al. 1983; Shawky et al. 2011). This variation in consanguinity rates affects the burden of runs of homozygosity (ROH) and recessive disease risk, but this has not been systematically characterized at the national genomic level.

Recent population genomic initiatives have begun addressing representation gaps in the Middle East and North Africa. The Qatar Genome Program sequenced over 6,000 Qatari genomes, revealing distinct genetic substructure within Arabian Peninsula populations (Razali et al. 2021; Scott et al. 2016). The Moroccan Genome Project characterized 109 individuals and introduced a population-specific reference genome, which reduced variant-calling variability by approximately 45% compared to GRCh38 (El Fahime et al. 2025). The UAE-based Arab Pangenome Reference provided the first pangenome for the region (Nassir et al. 2025). In Africa, the H3Africa consortium has supported genomic studies across multiple countries, and the African Genome Variation Project has characterized genetic diversity across sub-Saharan populations (H3Africa Consortium et al. 2014; Gurdasani et al. 2015).

The Egypt Genome Project addresses these gaps by conducting nationwide sampling (Elmonem et al. 2024; Amer et al. 2025). The current study characterizes genomic variation through wholegenome sequencing and elucidates population structure through analysis of autosomal, mitochondrial, and Y-chromosomal variation. Additionally, it examines ROH to reveal within-country heterogeneity with implications for recessive disease risk, characterizes Human Leukocyte Antigen (HLA) class I allele frequencies relevant to transplant medicine, and assesses the cross-population transferability of European-derived polygenic risk score thresholds.

## Materials and Methods

### Ethical Approval and Recruitment

The Central Institutional Review Board (IRB)-Supreme Council of University Hospitals approved this study (Reference: NO-b043(V-3)). All participants provided written informed consent in accordance with the Declaration of Helsinki and Egypt’s Law for Research and Clinical Trials No. 214/2020. Unique identifiers were used to protect participants’ privacy, and data were stored on password-protected servers with restricted access, in accordance with the Egyptian Data Protection Law No. 151/2020 and the General Data Protection Regulation guidelines.

In collaboration with the Egypt Center for Research and Regenerative Medicine (ECRRM), Cairo University, Ain Shams University, the National Research Centre, Alexandria University, the Magdi Yacoub Foundation, Shefaa Al-Orman Oncology Hospital, and Mansoura University, we recruited 1,135 Egyptian citizens between March 2022 and December 2024 meeting the following inclusion criteria: (1) Egyptian nationality with Egyptian-born parents; (2) age 18 years or older; (3) no first-degree familial relationships with other participants; and (4) absence of known genetic disorders or chronic diseases at enrollment. Participants were recruited through eight clinical and research centers across Egypt and originated from 21 of 27 governorates, spanning Greater Cairo (Cairo, Giza, Qalyubia), the Alexandria region (Alexandria, Beheira, Marsa Matruh), the Delta region (Dakahlia, Damietta, Gharbia, Kafr El Sheikh, Monufia), the Canal region (Ismailia, Sharqia [recruitment site in eastern Delta], Suez), the Northern region of Upper Egypt (Beni Suef, Faiyum, Minya), the Central region of Upper Egypt (Asyut), and the Southern region of Upper Egypt (Luxor, Qena, Sohag).

### Sample Processing and Sequencing

Whole blood samples (10 mL) were collected in PAXgene Blood DNA Tubes (PreAnalytiX, Switzerland) by trained phlebotomists using standard venipuncture procedures. Samples were transported to the ECRRM central facility within 72 hours under controlled temperature conditions (2–8 ^∘^C). Upon receipt, samples were logged into a Laboratory Information Management System with unique barcodes and stored at −80 ^∘^C until DNA extraction.

DNA was extracted using the automated Chemagic 360-D Instrument (Revvity, USA) following the Chemagic DNA Blood 400-360 H96 protocol. Extracted DNA was quantified using a Qubit 4 Fluorometer (Thermo Fisher Scientific, USA). DNA purity was measured using a NanoDrop One spectrophotometer (Thermo Fisher Scientific, USA) by calculating the 260/280 and 260/230 ratios. Samples with a Genomic Quality Score of 3 or higher were selected for library preparation. DNA libraries were prepared using the TruSeq DNA PCR-free library preparation kit (Illumina, USA). Sequencing was performed on Illumina NovaSeq 6000 at 2 x 150 bp read length using NovaSeq 6000 S4 Sequencing Reagent Kit v1.5 (300 cycles).

### Variant Calling and Quality Control

The DRAGEN DNA pipeline (Illumina) performed primary analysis with default parameters. Samples sequenced before mid-2023 were processed with DRAGEN v3.9.5; subsequent samples used v4.0.3. Both versions employ the same core alignment and variant-calling algorithms, and no batch effects attributable to pipeline version were detected in quality metrics. BCL files were converted to FASTQ files; paired FASTQ files were mapped to the human reference genome GRCh38, generating CRAM files, which were used to produce VCF files.

For quality control, samples were assessed for read quality (Q) and mean sequencing coverage; those with Q < 30 or mean coverage < 20x were excluded. Sex discrepancies between samples and metadata were assessed using Sambamba (Tarasov et al. 2015). Relatedness was evaluated using VCFtools v0.1.15 with the relatedness2 flag, restricting analysis to individuals with a kinship coefficient < 0.25 (Danecek et al. 2011). GATK Variant Quality Score Recalibration filtered SNV sites at 99.5% sensitivity and indel sites at 95% sensitivity (Van der Auwera et al. 2014). Variants passing these filters were further restricted to high-confidence genomic regions defined by the Genome in a Bottle (GIAB) consortium for the HG001 reference sample, using BCFtools v1.18 (Danecek et al. 2021); variants outside these regions were excluded to minimize false positives in difficult-to-map genomic contexts. Sensitivity and precision were evaluated against the GIAB HG001 truth set within these high-confidence regions; DRAGEN v3.9-v4.0 achieves SNV F1 scores exceeding 0.999 and indel F1 scores exceeding 0.995 on this benchmark according to manufacturer documentation. The transition/transversion (Ti/Tv) ratio was computed using BCFtools stats as a standard quality metric for variant calls. Functional annotation was performed using ANNOVAR (Wang, Li, and Hakonarson 2010).

### Population Structure Analysis

Egyptian genetic data were compared to populations from the 1000 Genomes Project (1000 Genomes Project Consortium et al. 2015), including 671 Africans, 340 Admixed Americans, 466 East Asians, 520 Europeans, and 492 South Asians, as well as 298 Middle Easterners from the Human Genome Diversity Project (161 individuals) and the Arab Population Genetics Project (137 individuals). Egyptian and reference genotype data were merged using PLINK v2.0 on the intersection of shared variant positions. The merged dataset was filtered to retain autosomal biallelic SNPs with a minor allele frequency (MAF) of 0.01 or greater, a call rate of 90% or greater, and Hardy-Weinberg equilibrium *p* > 1 × 10^−6^, using PLINK v2.0 (Chang et al. 2015).

Per-population allele frequencies were computed from the merged VCF using BCFtools for seven superpopulations (Egyptian, Middle Eastern, European, African, East Asian, South Asian, Admixed American). The degree of association between Egyptians and other populations was examined using Pearson correlation coefficients and linear regression. To ensure fair comparisons across all populations, correlations were computed on a common intersection of biallelic SNPs with non-missing allele frequency data in all seven populations and MAF ≥ 0.01 in both the Egyptian and reference population. Variants with absolute allele frequency differences exceeding 0.3 were classified as divergent. Shared variant counts per pairwise comparison and the number of divergent variants are reported.

Principal component analysis was performed using PLINK v2.0 (Chang et al. 2015) on genomewide autosomal biallelic SNPs retained after quality control (minor allele frequency ≥ 0.01, call rate ≥ 90%, Hardy-Weinberg equilibrium *p* > 1 × 10^−6^) and linkage disequilibrium (LD) pruning (window size 50 SNPs, step size 5, *τ*^2^< 0.2). ADMIXTURE v1.3.0 (Alexander, Novembre, and Lange 2009) estimated ancestral structure using the same LD-pruned SNP set, with 5-fold cross-validation over K values from 2 to 12; the optimal K was selected as the value minimizing cross-validation error. Runs of homozygosity were identified using PLINK v1.9 with the –homozyg flag (minimum 100 SNPs per segment, minimum segment length 1,000 kb, maximum 1 heterozygote per window, scanning window of 50 SNPs). These parameters follow established conventions for WGS-based ROH detection (Ceballos et al. 2018), and results were qualitatively stable when the minimum segment length was varied between 500 kb and 1,500 kb (data not shown). Genetic differentiation (fixation index, *F_ST_*) was evaluated using PLINK v2.0. Differences in ROH burden across populations and Egyptian subregions were assessed using the Kruskal-Wallis test, followed by pairwise Wilcoxon rank-sum tests with Benjamini-Hochberg (BH) correction for multiple comparisons. The association between self-reported parental consanguinity and total ROH burden was assessed using the Wilcoxon rank-sum test, with effect size quantified by Cliff’s delta.

### Uniparental Marker Analysis

Y-chromosome haplogroup classification was performed using HaploGrouper (Jagadeesan et al. 2021) and compared each male to the International Society of Genetic Genealogy phylogenetic tree. Mitochondrial haplogroups were assigned using Haplogrep 3 (Schönherr et al. 2023) with PhyloTree 17 (Oven 2015).

### HLA Typing

The DRAGEN HLA Caller (versions 3.9.5 and 4.0.3) was used to type Human Leukocyte Antigen (HLA) class I alleles (HLA-A, HLA-B, and HLA-C) within the Major Histocompatibility Complex. Reads mapping to HLA loci were aligned against a reference database of known HLA allele sequences, and the most probable allele pair for each locus was determined using integer linear programming in conjunction with population-level HLA allele frequencies.

### Polygenic Risk Score Analysis

Polygenic risk scores (PRS) were computed for each individual using pre-computed effect sizes from the PGS Catalog (Lambert et al. 2021). Scores were calculated for conditions with a minimum of 30% variant coverage in the merged dataset, yielding 10 assessable traits. Raw scores were standardized to z-scores using the global mean and standard deviation across all populations. To assess threshold transferability, the European population-specific 90th percentile z-score was computed, and the proportion of Egyptian individuals exceeding this threshold was computed empirically from the observed score distributions. Cohen’s d was computed as the standardized mean difference between Egyptian and European z-score distributions using the pooled standard deviation. Conditions exhibiting persistent bimodal distributions in the Egyptian cohort (type 2 diabetes PGS_000001, PGS_000014) were excluded, as were conditions falling below the 30% variant coverage threshold (coronary artery disease/hypertension PGS_000018).

### Clinical Variant Analysis

Genomic positions overlapping with ClinVar (version 2024-11) were identified using BCFtools isec. Only variants classified as Pathogenic or Likely Pathogenic according to American College of Medical Genetics and Genomics/Association for Molecular Pathology (ACMG/AMP) guidelines were retained for carrier frequency analysis; variants classified as Benign, Likely Benign, Uncertain Significance, or associated with susceptibility, risk factors, or protective phenotypes were excluded. Variants with conflicting interpretations of pathogenicity were excluded unless the majority of submitters classified the variant as Pathogenic or Likely Pathogenic. The ClinVar review status (number of submitters and presence of expert review) for each retained variant is reported in the supplementary materials. Variants supported by a single submitter without expert review criteria were retained but should be interpreted with caution, as their clinical significance may be revised in future ClinVar releases; such variants were not used to support clinical interpretation in isolation. Penetrance estimates were not available for most variants and are not incorporated into the carrier frequency calculations; the reported frequencies therefore represent genetic carrier status rather than clinical disease risk. Genotype calls were subjected to quality control filters requiring mapping quality (MQ) ≥ 30, read depth (DP) ≥ 20, and genotype quality (GQ) ≥ 20. Genes with known pseudogene interference (*CYP21A2* and *HBA1*) were excluded to prevent alignment artifacts. Carrier frequencies were calculated as the proportion of heterozygous individuals among 1,024 samples, with exact binomial (Clopper-Pearson) 95% confidence intervals. Hardy-Weinberg equilibrium was assessed using exact tests (appropriate for rare variants with low expected genotype counts) with false discovery rate (FDR) correction. Regional heterogeneity in carrier frequencies across Egyptian subregions was assessed using Pearson *χ*^2^ tests on 2 x k contingency tables (carrier vs. non-carrier across k subregions); subregions with zero carriers were excluded from individual gene tests to maintain valid expected cell counts.

### Projected Affected Births

For each autosomal recessive gene, the expected number of affected births per year was estimated from the observed carrier frequency (*q*_het_) as follows. Under Hardy-Weinberg equilibrium, the allele frequency was derived as *q* = *q*_het_/2, and the expected proportion of affected (homozygous) births was computed as *q*^2^. To account for the elevated probability of homozygosity due to consanguinity, the adjusted proportion was computed as *q*^2^ + *Fq*(1 − *q*), where *F* is the inbreeding coefficient corresponding to the national consanguinity rate of 35.3% (Shawky et al. 2011). The inbreeding coefficient was approximated as *F* = *α*/16, where *α* is the proportion of consanguineous marriages, following established population genetics conventions for first-cousin unions, the predominant form of consanguinity in Egypt. Projected affected births were obtained by multiplying the expected proportion by the annual birth cohort of approximately 2.2 million (Central Agency for Public Mobilization and Statistics, 2023).

## Results

### Variant Discovery

Whole-genome sequencing was performed on 1,135 healthy Egyptians. After quality control (mean coverage below 20x exclusion, sex concordance checks, and relatedness filtering at kinship coefficient < 0.25), 1,024 unrelated individuals (580 males, 444 females) were retained. Staged variant filtering through GATK VQSR, restriction to GIAB HG001 high-confidence regions, and functional annotation produced a final set of 51,342,784 variants (46,064,181 SNVs and 5,278,603 indels). Sequencing achieved a mean coverage of 35.9x (SD = 9.7x) with a Ti/Tv ratio of 2.08 (Table S1; Figure S1).

Of variants carried by at least one Egyptian individual, 17,140,291 (33.4%) were absent from dbSNP build 156, demonstrating that Egyptian genetic variation remains poorly captured by existing databases. The site frequency spectrum exhibited the expected L-shaped distribution (Figure S2), with rare variants (alternate allele frequency below 5%) comprising the majority of the catalog. Detailed variant statistics, including functional annotation breakdowns, singleton counts, and persample metrics, are provided in Supplementary Note 1.

### Allele Frequency Comparisons

Pairwise allele frequency comparisons across 6,495,768 biallelic SNPs (MAF ≥ 0.01) showed the strongest concordance between Egyptians and Middle Eastern populations (*τ* = 0.977; linear regression slope = 1.003, intercept = 0.003), followed by European (*τ* = 0.958), Admixed American (*τ* = 0.931), South Asian (*τ* = 0.927), East Asian (*τ* = 0.825), and African (*τ* = 0.823) populations (Figure S3). The lower correlation with African populations likely reflects the absence of North and Northeast African reference datasets; all available African reference populations are sub-Saharan (Yoruba, Luhya, Esan, Gambian Mandinka, Mende). The observed correlation therefore reflects genetic distance from sub-Saharan Africa rather than from the African continent as a whole. Divergent variants (absolute allele frequency difference exceeding 0.3) were most numerous in East Asian comparisons (14,989 variants) and least in European (1,050) comparisons.

### Population Structure

Principal component analysis positioned Egyptians on the European-African axis, proximate to Middle Eastern (including North African) and European populations, distinguishing them from sub-Saharan Africans and East Asians (Figure 1). The first principal component explained 67.1% of the genetic variance, and the second explained 12.0%.

**Figure 1:**
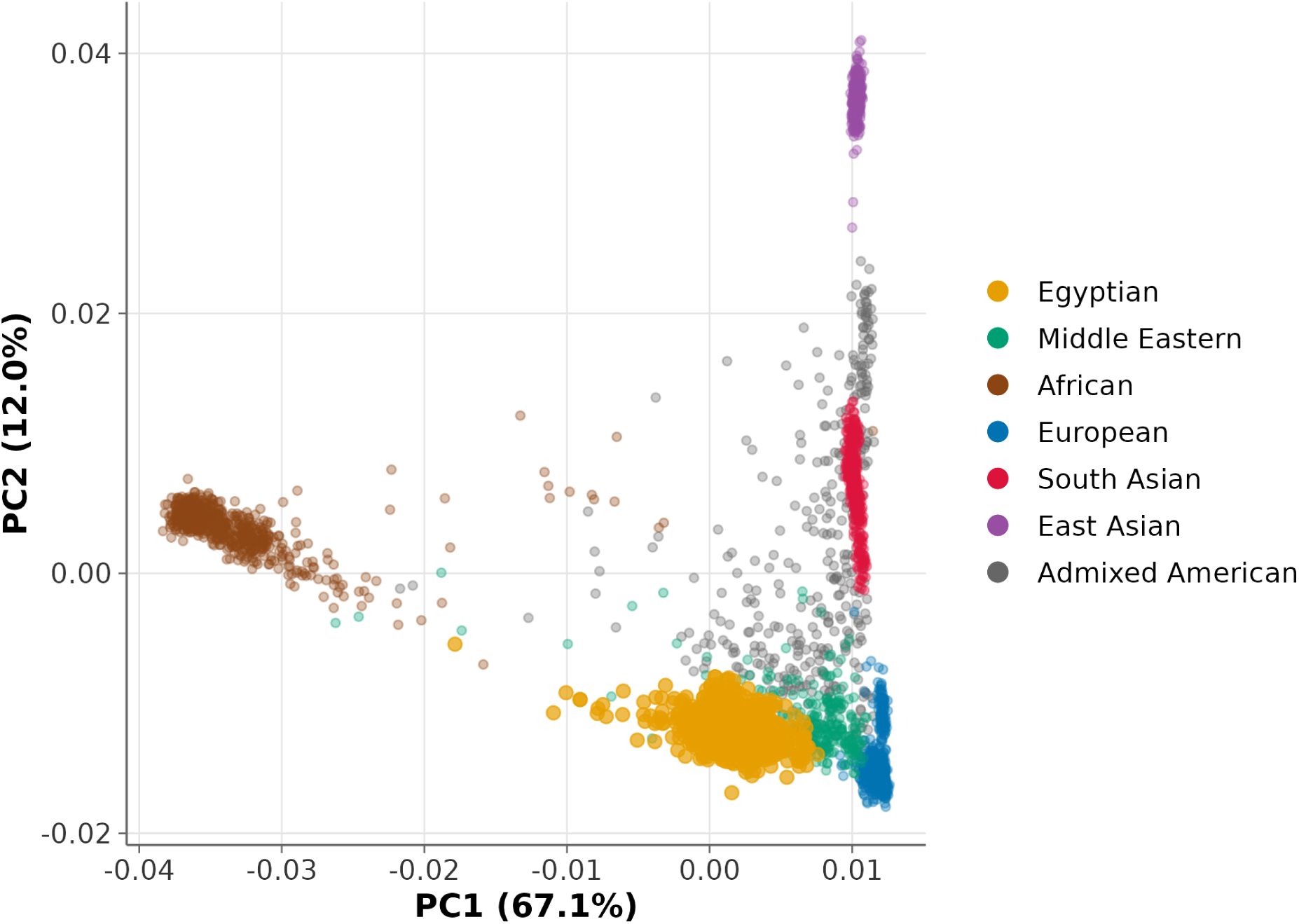
Principal component analysis of Egyptian (*n* = 1,024) and global reference populations (Middle Eastern, *n* = 298; African, *n* = 671; European, *n* = 520; South Asian, *n* = 492; East Asian, *n* = 466; Admixed American, *n* = 340). PC1 explains 67.1% of the genetic variance; PC2 explains 12.0%. Each point represents an individual, colored by superpopulation.

ADMIXTURE analysis at K = 7, the optimal value minimizing cross-validation error (Figure S4), revealed complex ancestry in the Egyptian population (Figure 2). Egyptians shared a dominant ancestry component with Middle Eastern populations (mean proportion 0.718 in Egyptians, 0.738 in Middle Easterners), reflecting deep shared West Eurasian ancestry. A smaller Egyptian-enriched component (mean 0.185 in Egyptians) distinguished them from neighboring populations. This component was highest in Egyptians, followed by Mozabite (0.165, *n* = 27) and Yemeni (0.069, *n* = 21) populations. Levantine and Gulf populations showed lower proportions: Bedouin (0.068, *n* = 46), Saudi (0.058, *n* = 28), Palestinian (0.044, *n* = 46), and Druze (0.001, *n* = 42). The remaining ancestry in Egyptians comprised European (0.048), African (0.035), South Asian (0.010), American-enriched (0.002), and East Asian (0.002) components (Figure S5).

**Figure 2:**
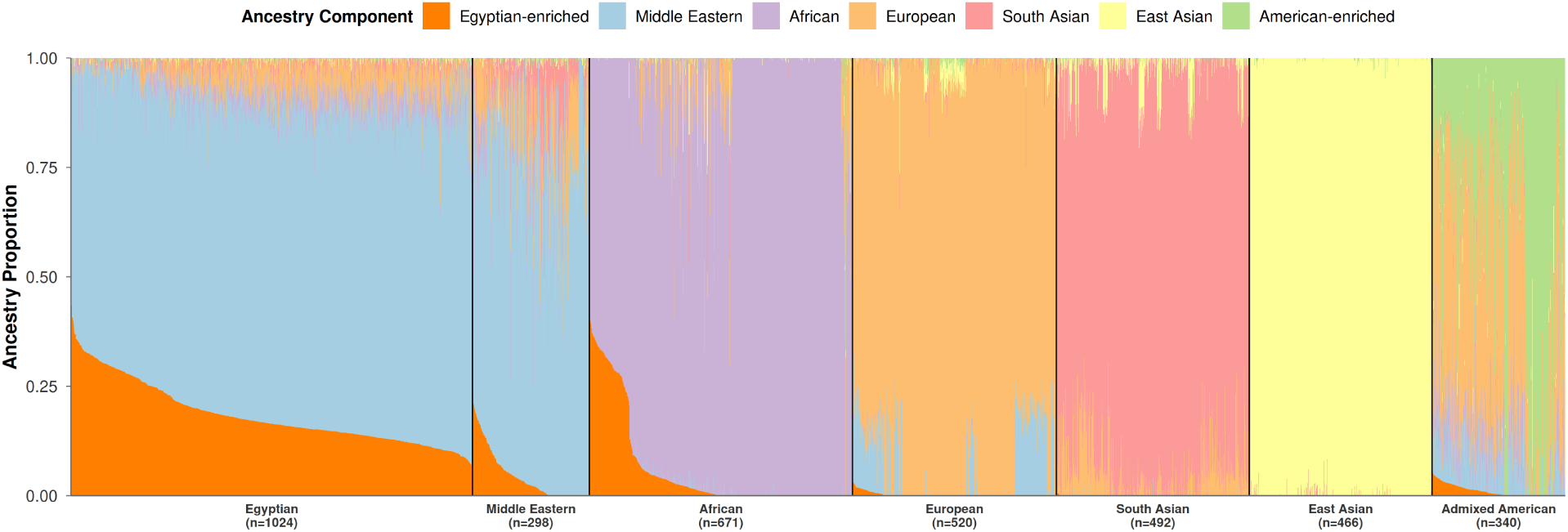
ADMIXTURE analysis of Egyptian and global reference populations (*n* = 3,811 total; sample sizes per population as in Figure 1). Ancestry proportions are shown at K = 7, the optimal value minimizing cross-validation error (Figure S4). Each vertical bar represents an individual; the y-axis shows the proportion of ancestry from each of seven inferred ancestral clusters. Populations are ordered by PCA-based Euclidean distance from Egyptians. The Egyptian-enriched component (18.5%) reflects population-specific allele frequency structure rather than a distinct ancestral source.

Weir-Cockerham *F_ST_* analysis on the LD-pruned SNP set quantified genetic differentiation between Egyptians and Middle Eastern populations (Figure S6). Yemeni (0.0021), Saudi (0.0027), and Bedouin (0.0029) populations showed the lowest *F_ST_* values. Levantine populations, including Palestinian (0.0031) and Syrian (0.0031), showed similarly low levels of differentiation. Iraqi (0.0034) and Emirati (0.0033) populations showed comparable values. Druze (0.0064) and Mozabite (0.0072) populations exhibited markedly higher *F_ST_* values. Jordan (*n*=3) and Oman (*n*=3) were excluded from quantitative *F_ST_* comparisons due to insufficient sample sizes; their negative *F_ST_* estimates (−0.0017 and −0.0007, respectively) are artifacts of sampling noise and are reported only for completeness in Figure S6. ADMIXTURE-based *F_ST_* estimates computed from mean ancestry proportions identified the same most-differentiated populations (Druze, Mozabite) and least-differentiated populations (Yemeni, Bedouin, Palestinian), though the two methods differed in the ranking of intermediate populations (Figure S7).

Subpopulation-level analysis using PCA-based Euclidean distance provided a granular view of genetic proximity across all 37 reference populations (Figure 3). Bedouin (0.014), Yemeni (0.016), and Saudi (0.016) were the closest populations to Egyptians, followed by Palestinian (0.018), Jordanian (0.019), and Syrian (0.021). Among non-Middle Eastern populations, Toscani (0.039) and Puerto Rican (0.040) were nearest, as expected given shared Mediterranean and admixed ancestry, respectively. Sub-Saharan African populations were the most distant, with Mende (0.096), Luhya (0.095), and Gambian Mandinka (0.091) showing the largest distances.

**Figure 3:**
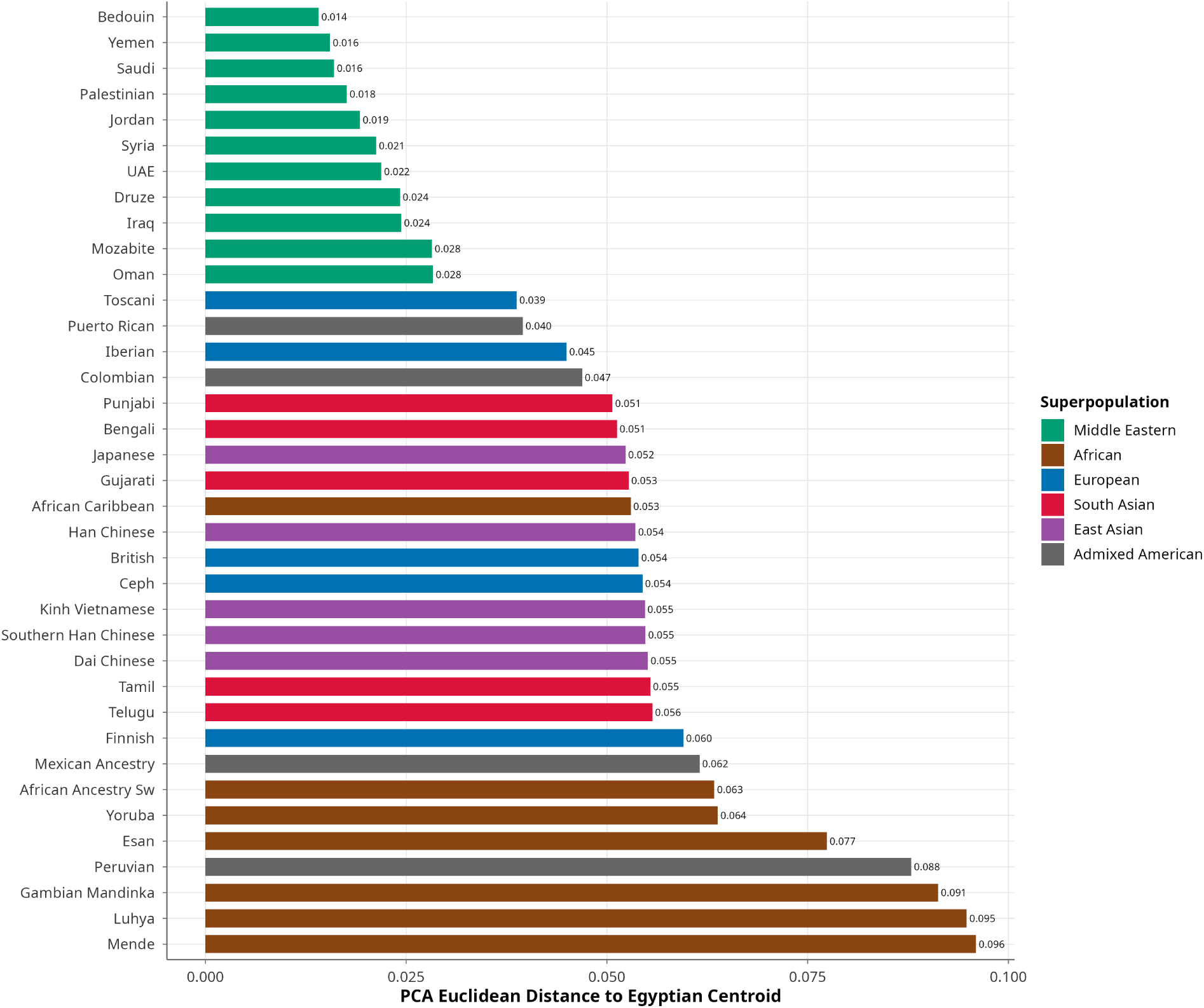
Genetic distance from Egyptians to 37 global reference populations. Euclidean distance was computed between each population’s centroid and the Egyptian centroid in genome-wide principal component space (PC1-PC20), providing a single quantitative measure of overall genetic differentiation. Populations are ordered from closest (top) to most distant (bottom). Colors indicate superpopulation grouping. Bedouin, Yemeni, and Saudi populations are the closest genetic neighbors of Egyptians.

### Runs of Homozygosity

ROH burden differed significantly across the seven population groups (Kruskal-Wallis *χ*^2^ = 757.67, *df* = 6, *p* < 2.2 × 10^−16^; Figure 4A). Egyptians (*n* = 1,024) exhibited a median total ROH length of 20,767 kb across a median of 108 segments, comparable to Admixed Americans (*n* = 340; median 26,550 kb) and East Asians (*n* = 466; 22,682 kb). Middle Eastern populations (*n* = 298) exhibited the highest median (68,924 kb), significantly greater than Egyptians (Wilcoxon *W* = 243,072, *p* < 2.2 × 10^−16^; Cliff’s delta = 0.73, 95% CI: 0.69-0.77). African (*n* = 671; 14,471 kb), European (*n* = 520; 18,666 kb), and South Asian (*n* = 492; 18,955 kb) populations showed lower median ROH burdens. Individual Egyptian ROH burdens ranged from near-zero to over 600,000 kb, indicating substantial within-population heterogeneity.

**Figure 4:**
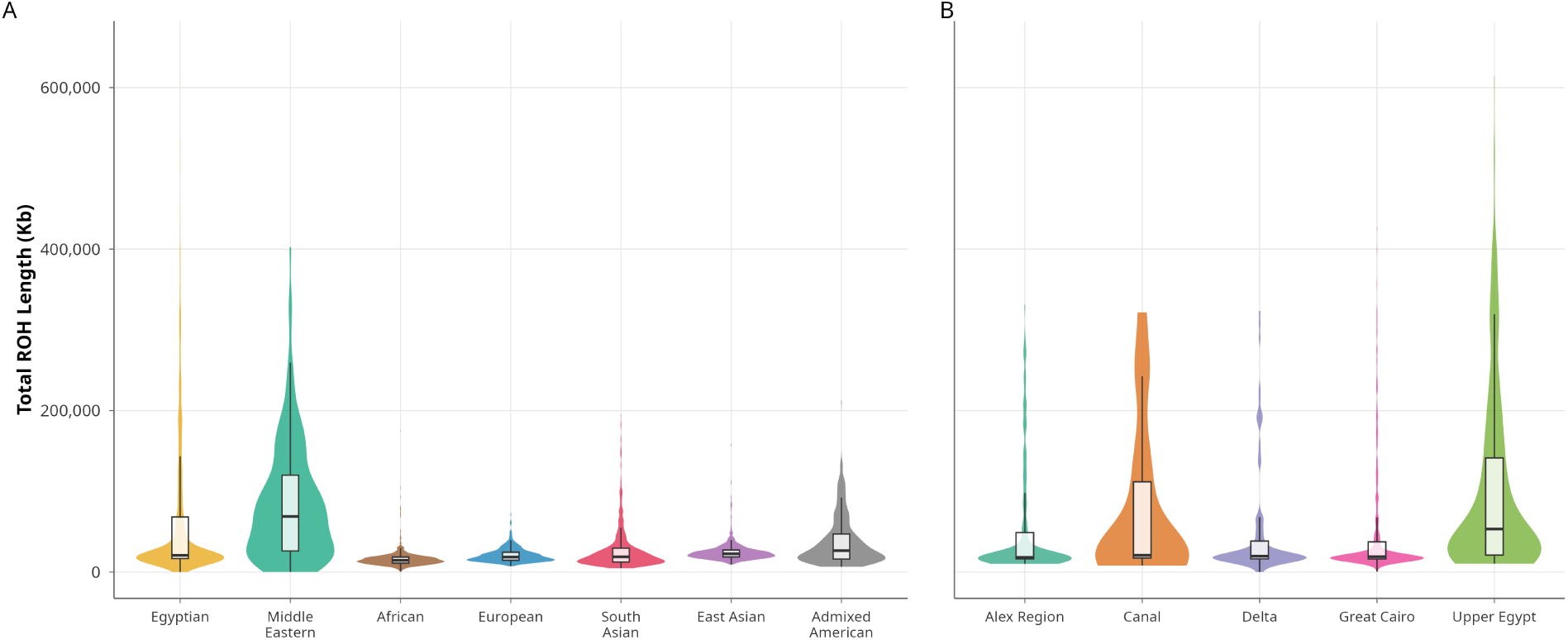
Runs of homozygosity in Egyptian and global populations. (A) Distribution of total ROH length (kilobases, reflecting recent consanguinity and population bottlenecks) across seven population groups (sample sizes as in Figure 1). Middle Eastern populations show the highest ROH burden, while Egyptians display substantial heterogeneity with a heavy right tail. (B) Distribution of total ROH length across five Egyptian subregions: Alex Region (*n* = 167), Canal (*n* = 20), Delta (*n* = 110), Greater Cairo (*n* = 450), and Upper Egypt (*n* = 277). Violin plots show density distributions; embedded boxplots indicate median and interquartile range.

This heterogeneity was geographically structured: ROH burden differed significantly across Egyptian subregions (Kruskal-Wallis *χ*^2^ = 105.34, *df* = 4, *p* < 2.2 × 10^−16^; Figure 4B). Upper Egypt (*n* = 277) showed the highest median (53,290 kb), significantly greater than Alex Region (*n* = 167; 18,295 kb; BH-adjusted *p* = 3.0 × 10^−12^), Delta (*n* = 110; 19,711 kb; *p* = 9.8 × 10^−10^), and Greater Cairo (*n* = 450; 19,064 kb; *p* < 2.0 × 10^−16^). Canal (*n* = 20; 20,780 kb) did not differ significantly from Upper Egypt (*p* = 0.21); however, with only 20 individuals, the Canal subregion is substantially underpowered for subgroup comparisons and these results should be interpreted with caution. The highest individual total ROH burden (613,967 kb) was observed in Upper Egypt. Among participants with available parental consanguinity data (*n* = 460), those reporting consanguineous parents (*n* = 98) had a median ROH burden of 157,894 kb compared to 18,412 kb for those with unrelated parents (*n* = 362), an 8.6-fold difference (Wilcoxon *W* = 32,480, *p* < 2.2 × 10^−16^; Cliff’s delta = 0.83, 95% CI: 0.77-0.88).

### Uniparental Markers

Y-chromosome and mitochondrial DNA lineages provided complementary evidence of dual ancestry (Figure 5). Paternal lineages were dominated by African haplogroup E (41.9%) and Middle Eastern haplogroup J (32.0%), with additional contributions from haplogroups R (9.1%), T (9.1%), and G (4.0%). Maternal lineages showed predominantly West Eurasian haplogroup H (47.3%) alongside African haplogroup L (27.1%), consistent with sex-biased admixture patterns previously observed in North African populations.

**Figure 5:**
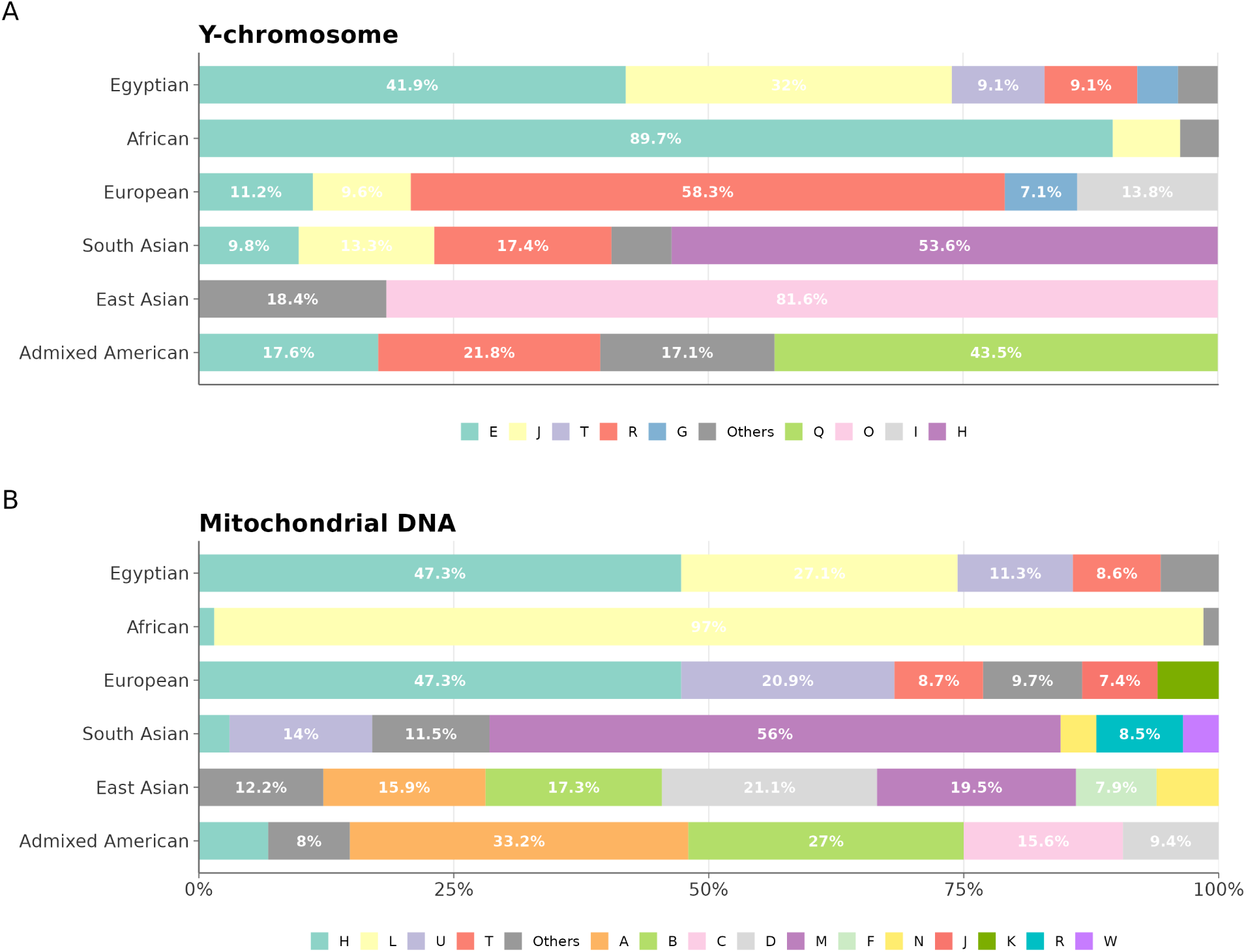
Uniparental haplogroup distributions across populations. (A) Y-chromosome haplogroups. The Egyptian population exhibits haplogroups E (41.9%), J (32.0%), R (9.1%), T (9.1%), and G (4.0%). (B) Mitochondrial DNA haplogroups. The Egyptian population is characterized by haplogroups H (47.3%), L (27.1%), U (11.3%), and T (8.6%).

### HLA Class I Frequencies

HLA class I typing identified A*02:01 (7.10%), B*41:01 (3.80%), and C*07:01 (6.90%) as the most frequent alleles at each locus, establishing baseline immunogenetic data for the Egyptian population (Figure 6). The elevated frequency of B*41:01 is a distinguishing feature of the Egyptian profile, as this allele is less common in European populations but has been reported at higher frequencies in other MENA groups (Arrieta-Bolanos, Hernandez-Zaragoza, and Barquera 2023). Detailed allele frequency distributions are provided in Supplementary Note 2.

**Figure 6:**
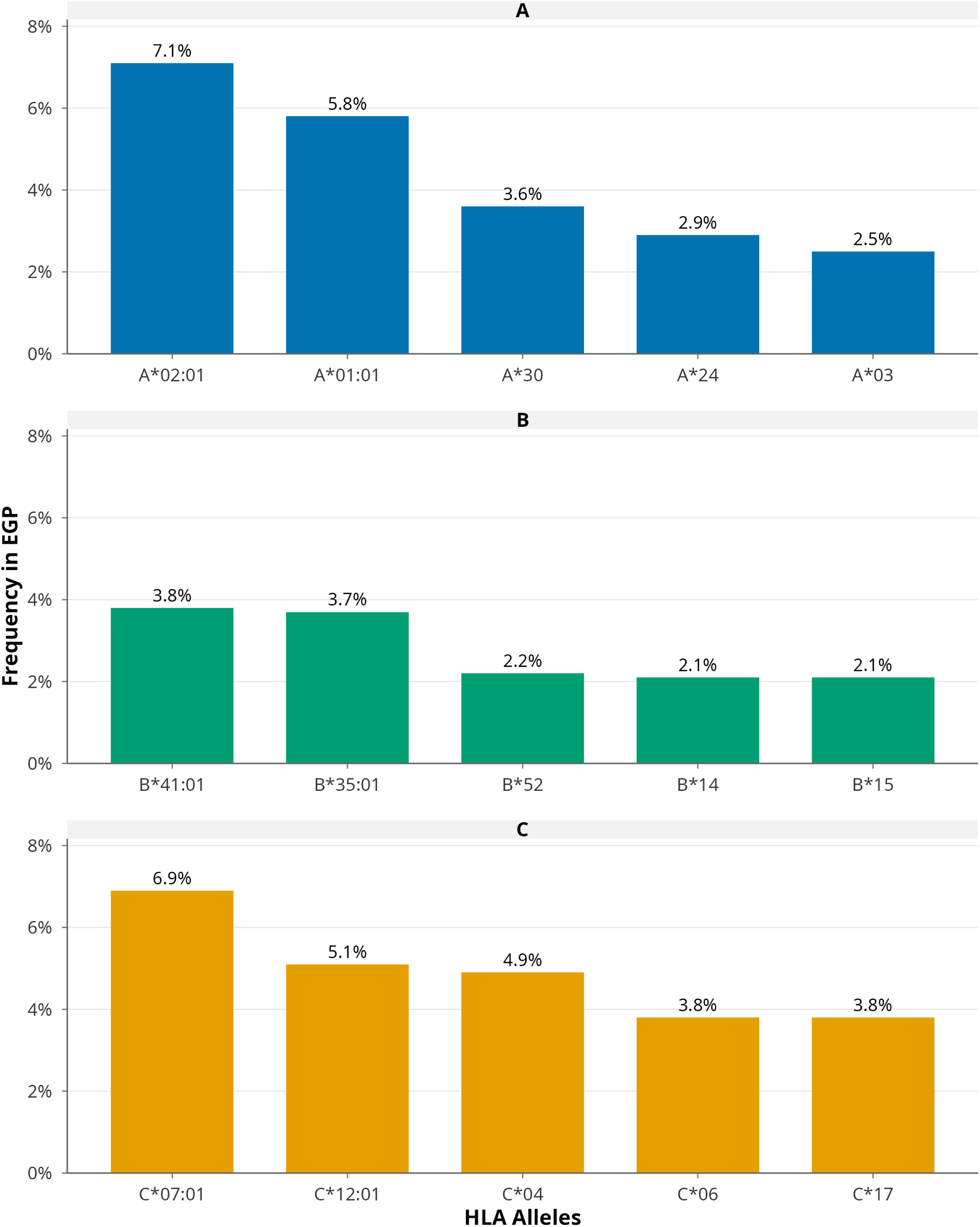
Most frequent HLA class I alleles in the Egyptian population (*n* = 1,024). Frequency distribution of the top five alleles at each HLA class I locus: (A) HLA-A, (B) HLA-B, (C) HLA-C. Allele frequencies represent the proportion of all typed alleles at each locus.

### Cross-Population Transferability of Polygenic Risk Scores

To assess the transferability of European-derived risk models to the Egyptian population, PRS were computed using effect sizes from the PGS Catalog and evaluated for conditions meeting a minimum variant coverage threshold of 30%, yielding 10 assessable traits. Scores were standardized as z-scores using the global mean and standard deviation across all populations. The European population-specific 90th percentile threshold was then applied, and the proportion of Egyptian individuals exceeding this threshold was computed empirically from the observed distributions.

Across several cardiometabolic traits, we observed substantial shifts in PRS distributions between Egyptian and European populations (Figure 7A). These distributional differences translated into marked deviations in threshold-based risk stratification. When the European-derived high-risk threshold (top 10%) was applied to the Egyptian cohort, 83.3% of individuals were classified as high risk for stroke (PGS_000027), 76.4% for chronic kidney disease (PGS_000016), and 72.8% for gout (PGS_000017), corresponding to approximately 7- to 8-fold inflation relative to the expected 10% (Figure 7B; Table 1). Middle Eastern reference populations also showed elevated proportions above the European threshold (43.0%, 51.7%, and 44.3% for the same traits, respectively), though to a lesser degree than Egyptians.

**Figure 7:**
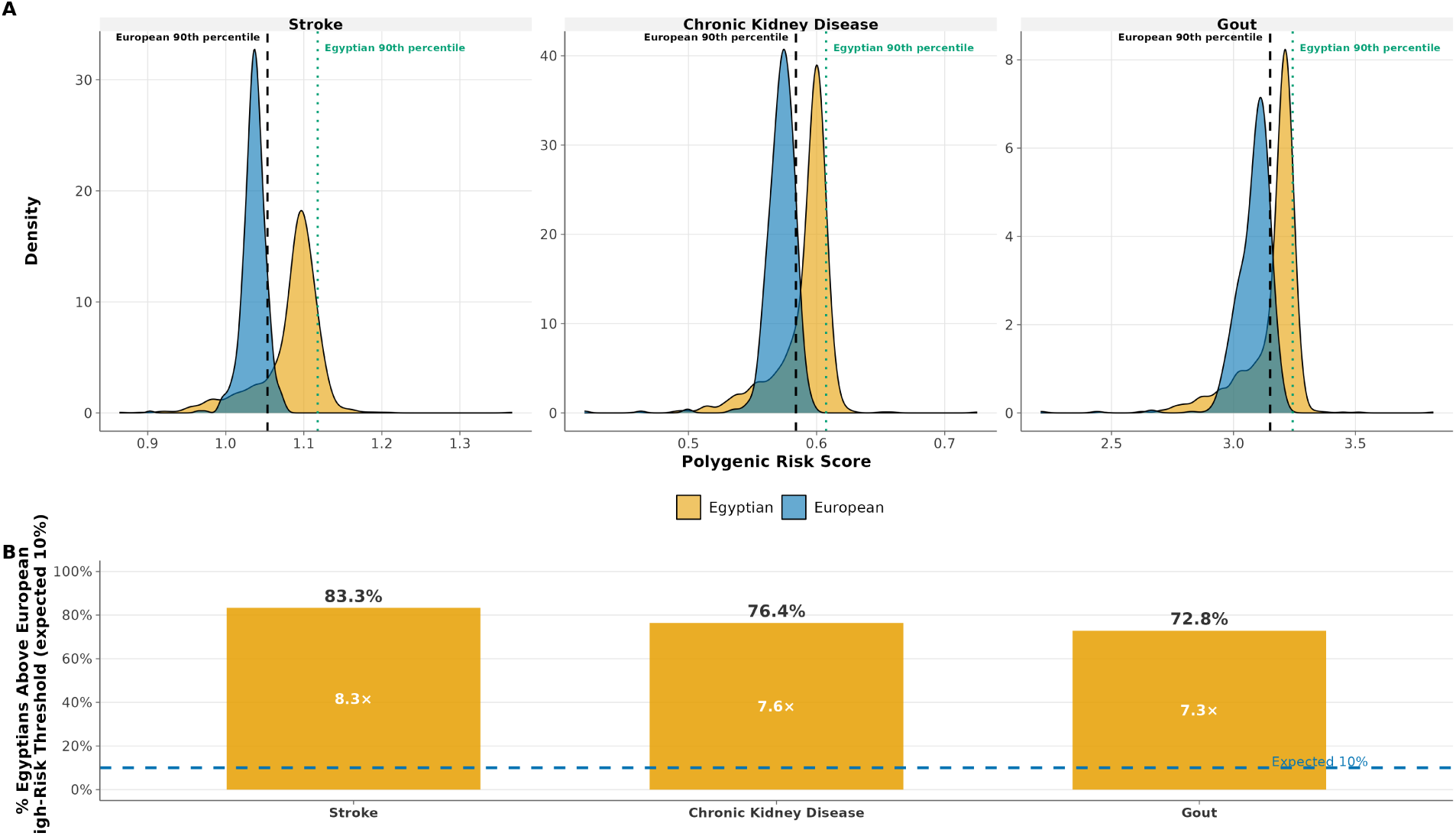
Cross-population differences in polygenic risk score distributions and threshold interpretation. (A) Density distributions of polygenic risk scores for stroke, chronic kidney disease, and gout in Egyptian and European populations. Vertical lines indicate the European 90th percentile (black dashed) and Egyptian 90th percentile (green dotted) thresholds. (B) Proportion of Egyptians exceeding the European high-risk threshold (top 10%). European-derived thresholds assign substantially higher proportions of Egyptians to the high-risk category (83.3%, 76.4%, and 72.8% for stroke, chronic kidney disease, and gout, respectively), corresponding to approximately 7- to 8-fold inflation relative to the expected 10%. These results indicate systematic differences in threshold calibration across populations.

**Table 1.** Cross-population transferability of polygenic risk scores: proportion of Egyptians exceeding European-derived high-risk thresholds.

| Trait | PGS ID | Cov. | EGY Z | EUR Z | d | EGY >P90 | ME >P90 |
| --- | --- | --- | --- | --- | --- | --- | --- |
| Stroke | PGS_000027 | 49.8 | 1.121 | -0.569 | 1.55 | 83.3% | 43.0% |
| CKD | PGS_000016 | 33.0 | 1.133 | -0.601 | 1.61 | 76.4% | 51.7% |
| Gout | PGS_000017 | 31.1 | 1.101 | -0.548 | 1.61 | 72.8% | 44.3% |

CKD = chronic kidney disease. Cov. = variant coverage, the proportion (%) of PGS Catalog score variants present in the merged dataset. EGY Z and EUR Z = mean z-scores for Egyptian and European populations, standardized using the global mean and standard deviation. d = Cohen’s d using pooled standard deviation. EGY >P90 and ME >P90 = proportion of Egyptian and Middle Eastern individuals above the European 90th percentile, computed empirically. Traits with bimodal distributions (type 2 diabetes) or variant coverage below 30% (coronary artery disease/hypertension PGS\_000018) were excluded.

These results indicate systematic differences in PRS distribution and threshold calibration across populations rather than true differences in underlying disease burden, consistent with the effect size differences observed between populations (Cohen’s d = 1.55-1.61). The observed shifts reflect known limitations in PRS portability, where differences in allele frequencies, LD structure, and effect size estimation contribute to reduced transferability across ancestries (Martin et al. 2019).

This effect was not uniform across traits. While cardiometabolic traits exhibited pronounced distributional shifts, other conditions showed closer alignment between populations. Rheumatoid arthritis and schizophrenia demonstrated near-overlapping PRS distributions and similar 90th percentile thresholds in Egyptian and European cohorts (Figure S8). In these cases, the proportion of Egyptian individuals exceeding the European high-risk threshold approximated the expected 10% (8.6% and 12.3%, respectively), indicating relatively preserved cross-population transferability.

PRS variant coverage ranged from 31.1% (gout) to 49.8% (stroke), and incomplete coverage may contribute to the observed distributional shifts independently of true population-level differences in genetic risk. However, across 38 traits with any variant representation in the PGS Catalog (including those below the 30% coverage threshold used for primary analysis), the correlation between variant coverage and the magnitude of the Egyptian-European distributional shift was weak and negative (*τ* = −0.23), indicating that lower coverage was not associated with larger shifts. Mean effect sizes were similar across coverage bins (|Cohen’s d|: 0.43 at <40% coverage, 0.28 at 40- 70%, 0.21 at >70%), suggesting that the large cardiometabolic shifts reflect trait-specific genetic architecture rather than coverage artifacts. Several conditions of clinical relevance were excluded: coronary artery disease/hypertension (PGS_000018) fell below the 30% coverage threshold, and type 2 diabetes (PGS_000001, PGS_000014) exhibited persistent bimodal distributions in the Egyptian cohort that could not be resolved through population stratification or sex stratification. This bimodality may reflect population substructure within the Egyptian cohort, score construction artifacts arising from the European training data, or interactions between ancestry-specific LD patterns and the T2D genetic architecture; resolving its origin will require larger sample sizes and additional reference populations. These analyses were not validated against phenotypic outcomes; the observed threshold differences therefore reflect distributional shifts in genetic risk scores rather than differences in clinical disease prevalence or predictive accuracy. These findings demonstrate that PRS transferability is trait-dependent and reinforce the need for population-specific calibration when applying PRS-based risk stratification across diverse populations.

### Carrier Frequencies for Mendelian Disease Genes

Integration of EGP1K data with ClinVar (version 2024-11) identified 1,332 variants classified as Pathogenic or Likely Pathogenic across 352,577 clinically annotated positions (Figure S9). Carrier frequencies were estimated for 13 autosomal recessive Mendelian disease genes (Table 2; Figure 8).

**Figure 8:**
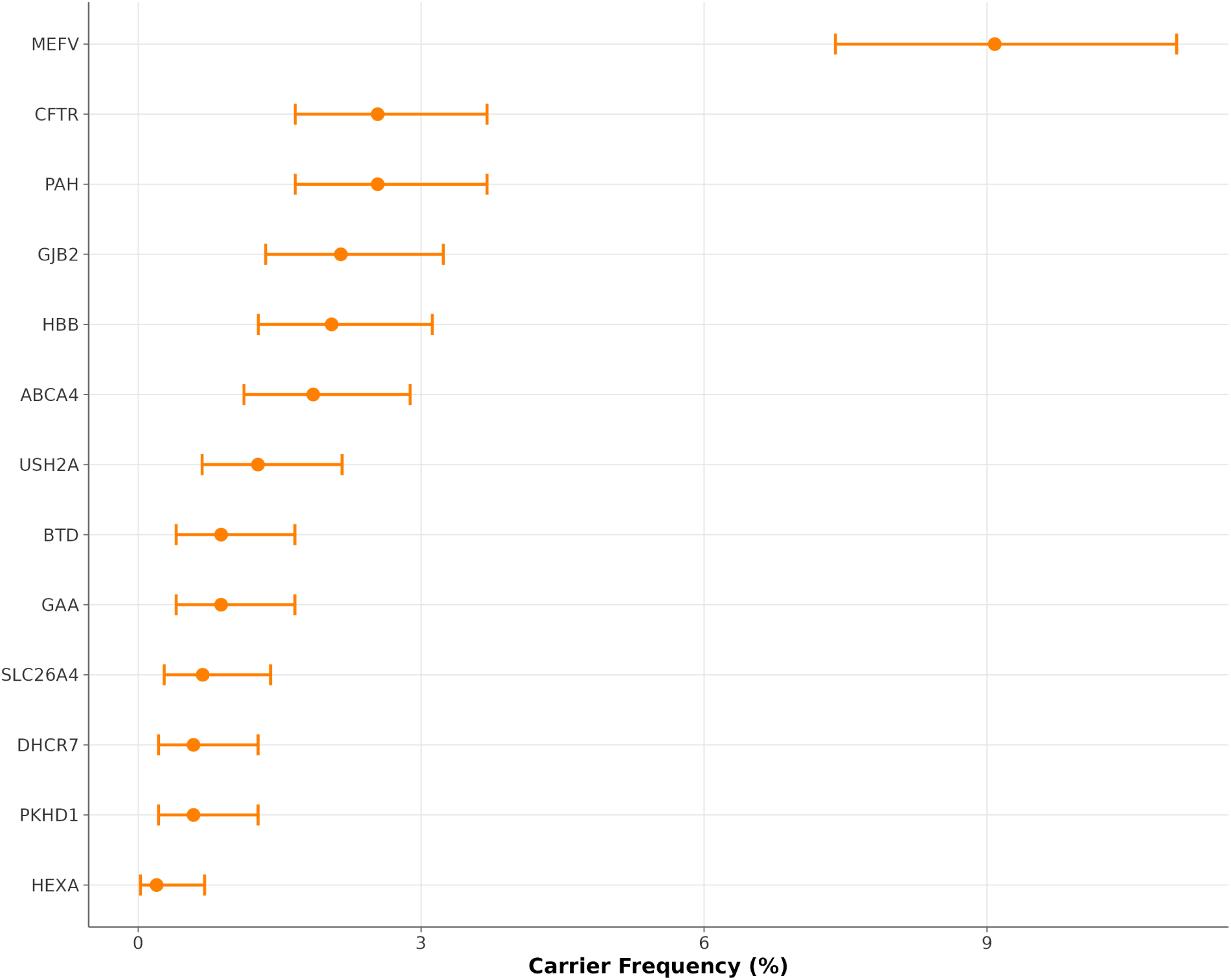
Carrier frequencies for autosomal recessive Mendelian disease genes in the Egyptian population (*n* = 1,024). Forest plot displaying heterozygous carrier frequencies with exact binomial (Clopper-Pearson) 95% confidence intervals for 13 autosomal recessive genes. Carrier frequency is defined as the proportion of individuals heterozygous for at least one pathogenic or likely pathogenic variant per gene. Variants were classified according to ClinVar (version 2024-11). *LDLR* (autosomal dominant) is reported separately in the main text.

**Table 2.**
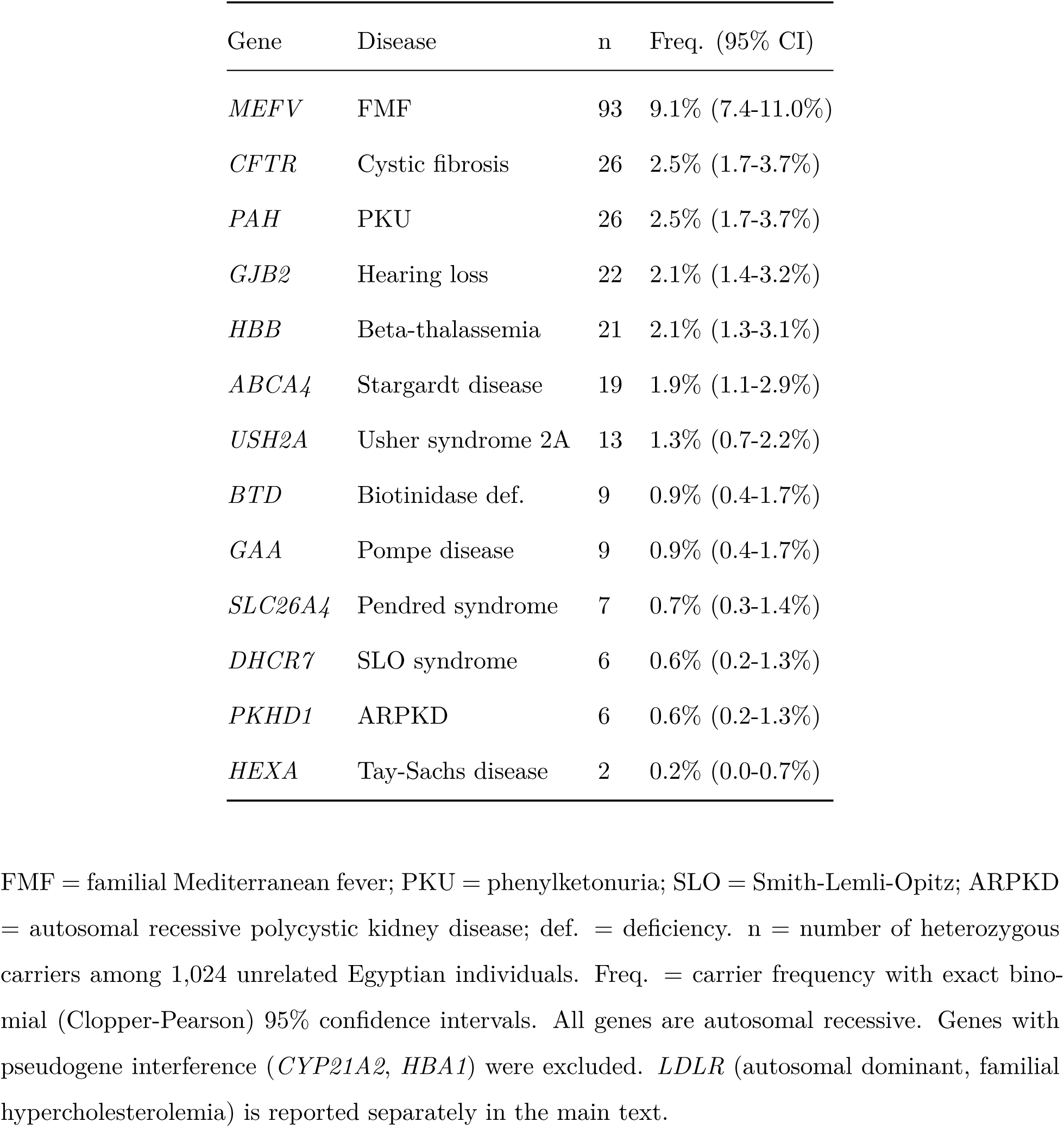
Carrier frequencies for Mendelian disease genes in the Egyptian population.

The highest carrier frequency was observed for *MEFV* (Familial Mediterranean Fever) at 9.1% (95% CI: 7.4-11.0%; 93 carriers across seven pathogenic variants; Table S2). This count includes two compound heterozygotes (Table S3) who are expected to be clinically affected. Additional genes with carrier frequencies exceeding 2% included *CFTR* (cystic fibrosis, 2.5%), *PAH* (phenylketonuria, 2.5%), *GJB2* (hereditary hearing loss, 2.1%), and *HBB* (beta-thalassemia, 2.1%). All tested variants were in Hardy-Weinberg equilibrium after FDR correction (Table S4). The distribution of rare pathogenic variants per individual is shown in Figure S10, and ClinVar evidence quality is shown in Figure S11. Comparison against published reference populations (Figure 9) showed that *MEFV* and *PAH* carrier frequencies in Egyptians exceed European estimates, while *CFTR* is lower than European pooled values (Table S5). When projected to Egypt’s annual birth cohort and adjusted for the national consanguinity rate of 35.3% (Shawky et al. 2011), *MEFV* alone accounts for an estimated 6,600 affected births per year, followed by *CFTR*, *PAH*, *GJB2*, and *HBB* (Figure 10). Separately, seven individuals (0.7%, 95% CI: 0.3-1.4%) were heterozygous for pathogenic *LDLR* variants associated with autosomal dominant Familial Hypercholesterolemia. Because *LDLR* variants are dominantly acting, these individuals are expected to be clinically affected rather than asymptomatic carriers, despite being recruited as healthy volunteers; this finding illustrates the limits of clinical screening in population recruitment and suggests that a fraction of apparently healthy participants may harbor undiagnosed monogenic conditions. Regional variation in carrier frequency was significant for *MEFV* (*χ*^2^ = 19.76, *df* = 4, *p* = 0.00056) and *GAA* (*χ*^2^ = 10.62, *df* = 3, *p* = 0.014).

**Figure 9:**
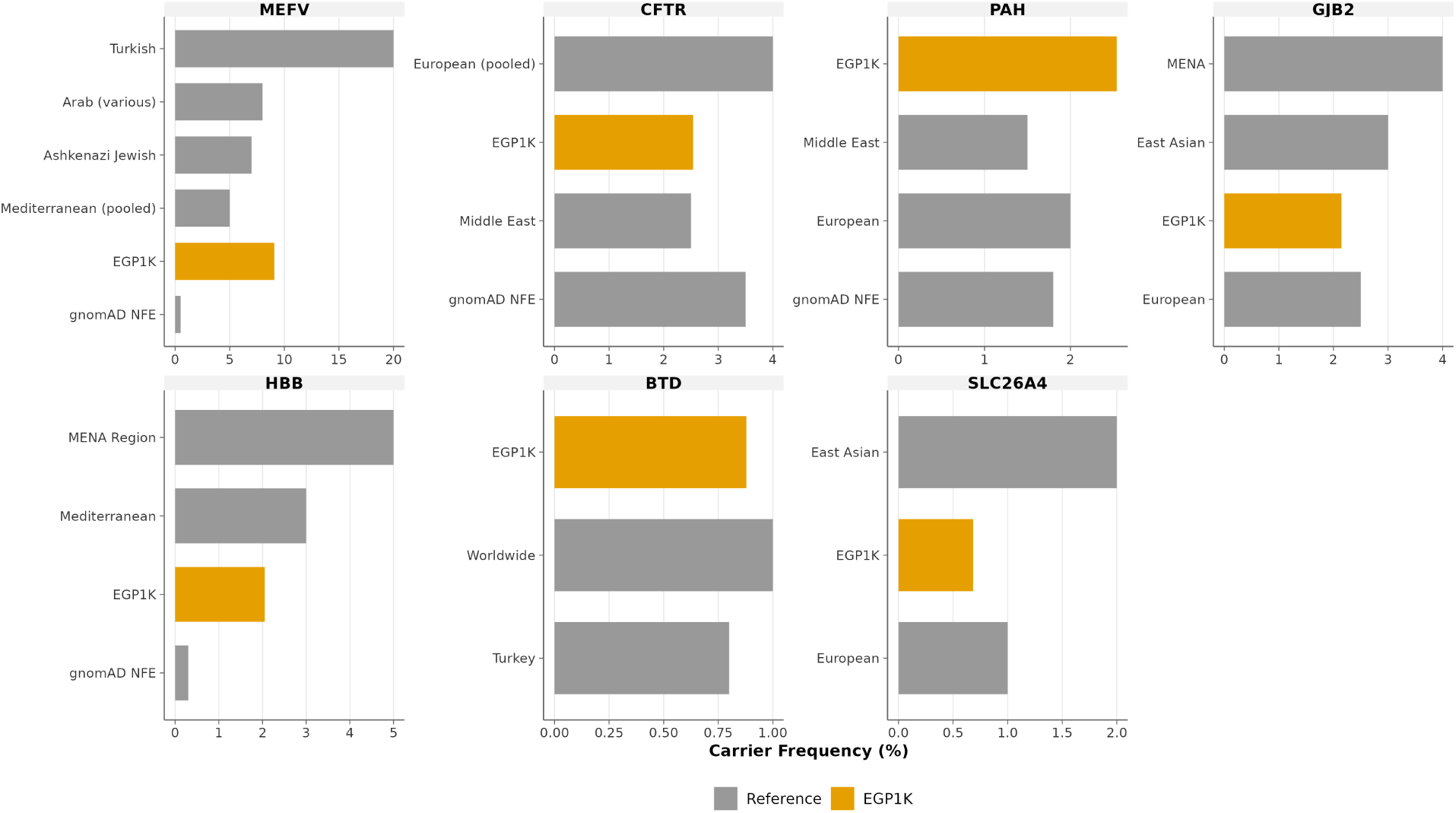
Comparison of Egyptian carrier frequencies with published reference populations. Grouped horizontal bar charts showing EGP1K carrier frequencies (amber bars, labeled “EGP1K”) alongside published reference values (gray bars) for seven genes with available comparison data (Table S5). Reference populations vary by gene based on available literature. Genes without published comparisons are shown in Figure 8.

**Figure 10:**
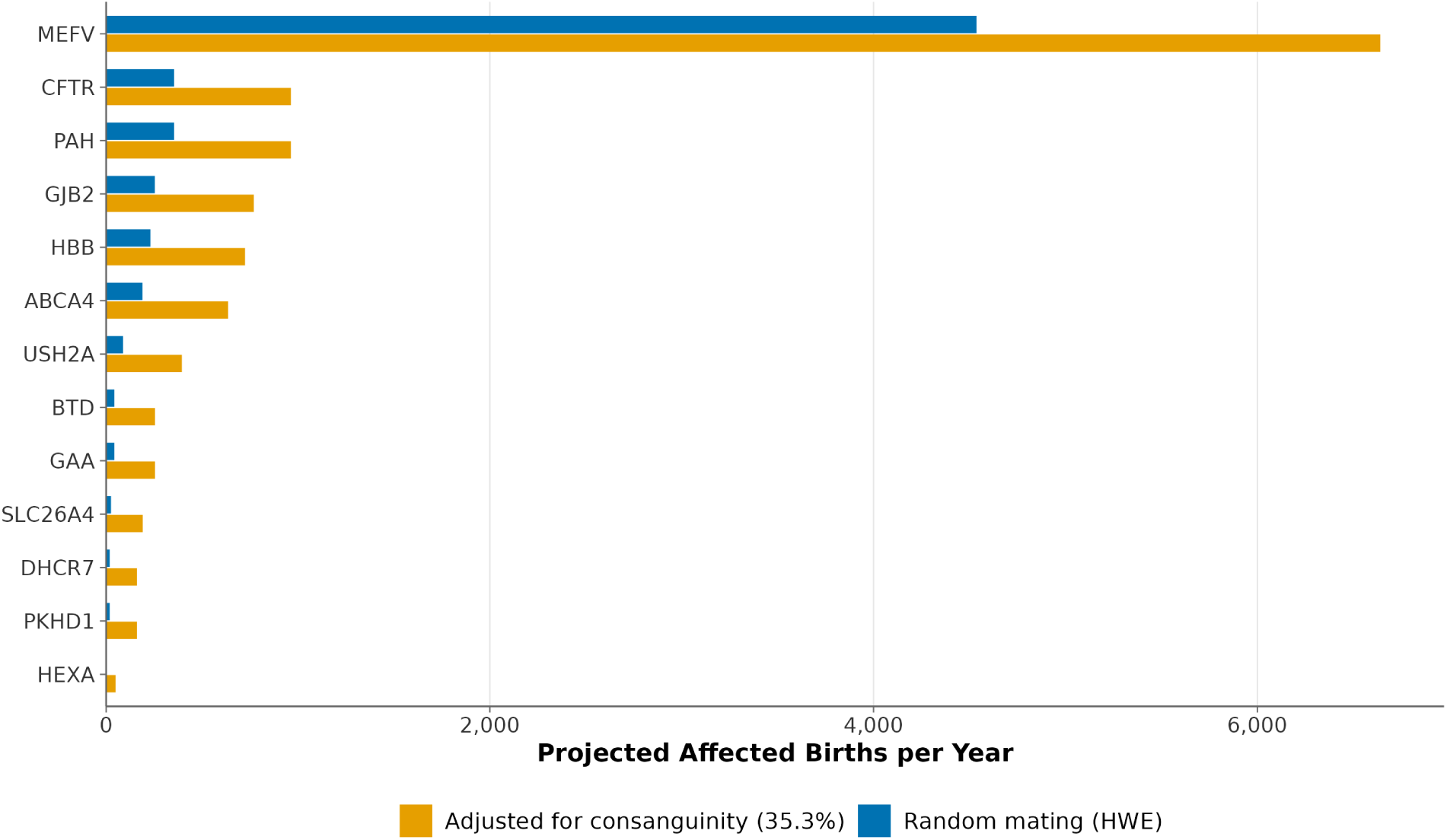
Projected affected births per year for autosomal recessive Mendelian diseases in Egypt. Horizontal bar chart comparing projected affected births under Hardy-Weinberg equilibrium assumptions (blue) and after adjustment for the national consanguinity rate of 35.3% (Shawky et al. 2011) (amber), based on an annual birth cohort of approximately 2.2 million (CAPMAS, 2023). *MEFV* (Familial Mediterranean Fever) shows the highest projected burden under both models.

## Discussion

With 1,024 individuals from 21 governorates and over 51 million variants cataloged, the Egypt Genome Project represents a nearly tenfold expansion over the previous 110-sample Egyptian reference (Wohlers et al. 2020). The fraction of novel variants (33.4% against dbSNP build 156) exceeds the 16.1% reported for a Han Chinese cohort of comparable size and sequencing depth (Juang, Lu, et al. 2020), and the 5.9% observed in the expanded 1000 Genomes Project high-coverage resource (Byrska-Bishop et al. 2022). This disparity reflects the degree to which Egyptian and MENA populations remain underrepresented in variant databases relative to East Asian and globally sampled cohorts.

### Population Structure and Genetic Affinity

Across all metrics examined, Egyptians show the strongest genetic affinity to Middle Eastern populations. The high allele frequency correlation (*τ* = 0.977) and near-unity regression slope (1.003) with Middle Eastern populations indicate strong concordance of common variant frequencies, a pattern expected given shared deep West Eurasian ancestry. The relatively low correlation with African reference populations, despite Egypt’s geographic position on the continent, arises from a known limitation of current reference panels: all available African references are sub-Saharan. Future comparisons against the Genome Aggregation Database (gnomAD) (Karczewski et al. 2020), which includes Middle Eastern samples, and the inclusion of North and Northeast African populations (e.g., Sudanese, Ethiopian, Somali) would provide a more complete picture of Egyptian genetic relationships within Africa. Principal component analysis and subpopulation-level distance analysis consistently place Egyptians alongside Middle Eastern and North African populations, with Bedouin, Yemeni, and Saudi as the closest genetic neighbors, in keeping with Egypt’s role as a bidirectional corridor for human migration (Serra-Vidal et al. 2019).

The subpopulation-level distance analysis (Figure 3) provides additional resolution. Among Middle Eastern populations, Bedouin and Yemeni are genetically closest to Egyptians, which is expected given the geographic continuity between the Nile Valley, the Sinai Peninsula, and the western Arabian Peninsula. The greater distance of Druze and Mozabite from Egyptians is attributable to their documented histories of genetic isolation (Shlush et al. 2008; Henn et al. 2012). Among non-Middle Eastern populations, Toscani (Southern Italian) are the nearest European group, in line with the long history of bidirectional gene flow across the Mediterranean basin linking North Africa and Southern Europe (Botigué et al. 2013). The admixed composition of Puerto Rican and Colombian populations, which include both European and African ancestral components, places them at intermediate distances. Sub-Saharan African populations show the greatest distances from Egyptians, with a gradient from admixed groups (African Caribbean, 0.053) to West African populations (Mende, 0.096), reflecting the deep divergence between North African and sub-Saharan genetic lineages (Henn et al. 2012).

### Ancestry Components

The ADMIXTURE results at K = 7 place Egyptians squarely within the West Eurasian genetic sphere, with 71.8% of ancestry shared with Middle Eastern populations. The Egyptian-enriched component, which distinguishes Egyptians from neighboring groups, is shared at appreciable levels only with Mozabite Berbers and at much lower proportions in Arabian Peninsula populations. This gradient suggests that the component captures ancestry specific to North African populations, likely tracing to autochthonous North African lineages rather than recent gene flow. The high proportion shared between Egyptians and Mozabites supports this interpretation, as both represent indigenous North African populations. Arabian Peninsula populations retain substantial ancestry from Natufian-related hunter-gatherers who inhabited both the Levant and Arabia during the Late Pleistocene and early Holocene (Lazaridis et al. 2016; Schuenemann et al. 2017), but this shared Natufian ancestry is captured by the dominant Middle Eastern component rather than the Egyptian-enriched one (Almarri et al. 2021; Rodriguez-Flores et al. 2016; Haber et al. 2017).

Weir-Cockerham *F_ST_* -based genetic distances corroborate the findings on ancestry proportions. The elevated differentiation observed for Druze (0.0064) and Mozabite (0.0072) populations aligns with their documented genetic isolation. Druze practice religious endogamy and the prohibition of conversion, which has maintained genetic differentiation from neighboring populations since the 11th century (Shlush et al. 2008). Mozabites represent an autochthonous North African Berber population that has experienced periods of genetic isolation and drift (Figure S7).

ADMIXTURE-derived ancestry proportions are sensitive to the composition of the reference panel. When Middle Eastern subpopulations (Bedouin, Yemeni, Saudi, Palestinian) are absent from the reference set, the North African component tends to absorb shared West Eurasian ancestry, inflating its apparent proportion. With the expanded reference panel used here, which includes 298 Middle Eastern individuals from 10 subpopulations, this shared ancestry is resolved as a distinct Middle Eastern component (71.8%), while the Egyptian-enriched component (18.5%) captures autochthonous North African ancestry. Analyses using fewer Middle Eastern references may therefore characterize the same underlying genetic structure differently, and apparent contradictions between studies should be interpreted in light of reference panel composition.

### Runs of Homozygosity

ROH analysis reveals population substructure with implications for medical genetics. Despite sharing substantial ancestry with Middle Eastern populations, Egyptians carry a markedly lower median ROH burden, with a large effect size (Cliff’s delta = 0.73). However, the wide range of individual values, extending from near-zero to over 600 Mb, indicates considerable within-population heterogeneity. The most extreme outlier, with over 613 Mb in ROH (approximately 20% of the genome), likely represents a product of close consanguinity.

Regional analysis localized this heterogeneity: Upper Egypt showed the highest median ROH length and harbored individuals with the longest ROH segments observed in this study. Extended ROH segments are associated with increased risk for recessive Mendelian disorders (Ceballos et al. 2018; Woods et al. 2006; Bittles and Black 2010; Leutenegger et al. 2011). Given documented consanguinity rates ranging from 17% in urban Egypt to 60% in rural Upper Egypt, these findings suggest that Upper Egypt may warrant targeted investigation of consanguinity practices and associated disease burden. Self-reported parental consanguinity was strongly associated with total ROH burden, with consanguineous individuals showing an 8.6-fold higher median ROH length than those with unrelated parents, consistent with the expected relationship between parental consanguinity and autozygosity.

### Uniparental Markers and HLA

The uniparental marker profiles reflect the dual African and West Eurasian heritage of Egyptians. The Y-chromosome profile positions Egyptians intermediate between African populations (where haplogroup E predominates) and Middle Eastern populations where haplogroup J is frequent. Haplogroup J has been associated with Neolithic demographic expansions from the Fertile Crescent (Arredi et al. 2004; Chiaroni et al. 2010; Semino et al. 2004; Di Giacomo et al. 2004), while haplogroup T (9.1%) is consistent with Levantine and Arabian Peninsula connections (Cadenas et al. 2008). Maternal lineages show predominantly West Eurasian haplogroup H (47.3%) alongside African haplogroup L (27.1%), contrasting with broader African populations where haplogroup L reaches 97% (Kujanová et al. 2009). HLA class I allele frequencies place Egyptians within the Levantine-Eastern Mediterranean cluster (Hajjej et al. 2018), with the elevated frequency of B*41:01 representing a distinctive feature of the Egyptian immunogenetic profile (Arrieta-Bolanos, Hernandez-Zaragoza, and Barquera 2023). These frequencies address the current absence of Egyptian representation in international HLA databases used for transplant matching and provide baseline data for investigating HLA-disease associations, particularly given Egypt’s historically high hepatitis C burden (Waked et al. 2014; Kouyoumjian, Chemaitelly, and Abu-Raddad 2018).

### PRS Transferability

The substantial cardiometabolic PRS distributional shifts reported above, with 73-83% of Egyptians exceeding the European 90th percentile, reflect differences in threshold calibration rather than true differences in underlying disease burden. These findings are consistent with prior work demonstrating reduced portability of PRS across ancestrally diverse populations (Martin et al. 2019; Ding et al. 2023).

At the same time, our results indicate that PRS transferability is not uniform across traits. Rheumatoid arthritis and schizophrenia showed relatively aligned PRS distributions and similar high-risk thresholds between populations, with proportions of individuals exceeding the European threshold approximating the expected 10% (Figure S8). This variability suggests that the degree of transferability depends on trait-specific genetic architecture, including factors such as polygenicity, effect size distribution, and differences in LD patterns across populations (Shim et al. 2023).

Given the analytical limitations detailed below, direct application of thresholds derived from European cohorts may lead to systematic misassignment of risk strata in underrepresented populations. Expanding genomic studies to include diverse populations and developing population-aware PRS models will be necessary for equitable implementation of genomic risk prediction.

### Carrier Frequencies and Clinical Implications

The 9.1% *MEFV* carrier frequency aligns with estimates from other Mediterranean and Middle Eastern populations (5-20% depending on ancestry; Aksentijevich et al. (1999); Yepiskoposyan and Harutyunyan (2007)) and has direct implications for genetic counseling given Egypt’s elevated consanguinity rates. Gene-by-gene comparison (Figure 9) shows that carrier screening programs calibrated to European populations would underestimate *MEFV* and *PAH* risk in Egyptian couples while providing reasonable estimates for *CFTR*. Given the significant regional variation observed for *MEFV* and *GAA* carrier frequencies, screening recommendations may need to account for geographic heterogeneity within Egypt. The carrier frequencies reported here are based exclusively on variants classified as Pathogenic or Likely Pathogenic in ClinVar, which is expected to sub-stantially underestimate the true carrier burden for many genes. Egyptians are likely to harbor population-specific disease-causing variants that have not yet been submitted to ClinVar, given that the majority of ClinVar submissions originate from European and other well-studied populations (Venner et al. 2024). Identifying and reporting these population-specific pathogenic variants is a key objective of the Egypt Genome Project, both to facilitate genetic counseling and family planning for affected families and to establish more accurate carrier frequency estimates for monogenic disease genes in the Egyptian population.

### Limitations

This study has several limitations. First, while participants originate from 21 of 27 governorates, six governorates are not represented (Aswan, New Valley, North Sinai, South Sinai, Port Said, and Red Sea), most of which are sparsely populated frontier regions; their absence may limit generalizability to those areas. Second, the healthy volunteer recruitment strategy may result in underestimation of carrier frequencies, particularly for conditions with clinically apparent heterozygote phenotypes. Third, *F_ST_* and ancestry estimates for populations with very small sample sizes (Jordan *n*=3, Oman *n*=3) should be interpreted with caution. Fourth, the African reference populations are exclusively sub-Saharan; the absence of North and Northeast African references limits interpretation of the Egyptian-African genetic relationship. Fifth, PRS variant coverage ranged from 31% to 50% for key traits, and no phenotypic outcome data were available for validation; the observed threshold differences therefore reflect distributional shifts rather than clinically validated risk predictions. Sixth, type 2 diabetes and coronary artery disease/hypertension PRS could not be included due to bimodal distributions and insufficient variant coverage, respectively, limiting the scope of the transferability analysis. Seventh, allele frequency comparisons were based on the 1000 Genomes Project and HGDP; comparison against gnomAD (Karczewski et al. 2020) is planned for future work. Eighth, carrier frequency estimates for genes with large allelic heterogeneity and limited expert panel coverage in ClinVar may be conservative; in particular, *PKHD1* (for which over 300 pathogenic variants have been reported but few have achieved multi-submitter or expert panel review status) yielded a carrier frequency of 0.6%, below the globally reported estimate of approximately 1 in 70 (Guay-Woodford and Desmond 2024), likely because genuine pathogenic variants supported only by single submitters were retained with caution but not used for primary carrier frequency estimation. Future reclassification of such variants by expert panels may revise these estimates upward.

## Conclusion

The Egypt Genome Project provides a large-scale genomic reference for Egypt, the most populous nation in the MENA region, with 1,024 individuals from 21 of 27 governorates and over 51 million variants cataloged.

Analysis of polygenic risk score distributions reveals substantial distributional miscalibration when European-derived thresholds are applied to Egyptians, with 73-83% exceeding the European 90th percentile for cardiometabolic traits. These distributional differences, observed despite incomplete variant coverage and without phenotypic validation, reinforce the case for developing population-specific calibration frameworks before PRS-based risk stratification is applied in clinical settings. Carrier frequency data provide reference values for population-appropriate screening programs. The regional heterogeneity in ROH burden, particularly in Upper Egypt, identifies priority areas for genetic counseling initiatives.

Ongoing work will expand the cohort and incorporate long-read sequencing for structural variant discovery. By characterizing genetic diversity in the most populous MENA nation, this work contributes to extending genomic resources to underrepresented MENA populations.

## Data Availability

Summary statistics, population-level allele frequencies, and allele frequency correlation data will be deposited in the European Variation Archive (EVA); accession numbers are pending. Individual-level genotypes are available through a controlled-access application to the Egypt Center for Research and Regenerative Medicine (ECRRM), with decisions made by the Egypt Genome Project Access Committee within 30 days of the application.

## Acknowledgments

We thank all participants who contributed samples to this study and the clinical staff at collaborating institutions who facilitated recruitment. This work was made possible by the Egypt Genome Project (EGP), funded by the Academy of Scientific Research and Technology (ASRT) and led by the Egypt Center for Research and Regenerative Medicine (ECRRM). We acknowledge the Central Agency for Public Mobilization and Statistics (CAPMAS) for designing the statistical sampling framework for the population genome project.

## Egypt Genome Consortium

The Egypt Genome Consortium coordinated recruitment, sample processing, sequencing, bioinformatics, and clinical interpretation across multiple Egyptian institutions. Members are listed below by working group.

**Egypt Genome Project (EGP) Lead Principal Investigator:** Khaled Amer (ECRRM; AFCM).

**Scientific Committee – Population Genome Working Group.** Ahmed Ihab (New Giza University); Ahmed Moustafa (ECRRM; AUC); Amal Mahmoud (NRC); Heba Hosni (MOHP); Heba Kassem (Alexandria University); Khaled Amer (ECRRM); Lamiaa Elwakil (MOHP); Mohamed Elhadidy (Nile University); Mohamed Salama (Mansoura University; AUC); Naglaa Kholousy (NRC); Rasha Elsherif (NCI); Sameh Saad (57357 Hospital); Sameh Soror (Helwan University); Samira Ezzat (Shefaa Al-Orman Hospital); Sherif El-Khamisy (University of Sheffield); Tarek Taha (ECRRM); Yasmine Aguib (Magdi Yacoub Heart Foundation); Yehia Z. Gad (Former Chair, NRC; NMEC).

**Scientific Committee.** Abdelrahman Zekri (NCI); Ahmed Ashour Ahmed (University of Oxford, United Kingdom); Amin Fouad (ECRRM); Azza Saleh Radwan (Theodore Bilharz Institute; Shefaa Al-Orman Hospital); Ezzat Elsobky (Ain Shams University); Gina El-Feky (ASRT); Heba Elsedafy (Ain Shams University); Hesham Ali Sadek (University of Texas Southwestern, USA); Hussein Khaled (NCI, Cairo University); Maged Mohamed El-Sadeq (ASRT); Maha El-Rabbat (Cairo University); Mahmoud Elhefnawy (NRC); Mohamed Awad Tag El-Din (Ain Shams University); Mohamed El Gohary (ECRRM); Mohamed Hassany (MOHP); Moustafa El Nakib (ECRRM); Neveen Nabil Abu El-Ainein (ASRT); Neveen Soliman (Chair, Cairo University); Sahar Kamal (Cairo University); Saleh Ibrahim (University of Luebeck, Germany); Solaf M. Elsayed (Ain Shams University).

**ECRRM Team. Project Management and Administration:** Moustafa El Nakib (CEO), Tarek El Nagdy. **Sample Preparation and Biobanking:** Gamal Okla, Hakam Hamed, Hashem

Ashour, Heba Hany, Rania Usama. **Central Genomic Lab:** Ahmed H. Badr, Fadya M. El-Garhy, Hasnaa El Shehaby, Marwa A. Zahra, May Amer, Nermeen T. Fahmy, Omnia M. Abdel-Haseeb, Tokka M. Hassan, Wael A. Hassan, Yara A. Daoud, Yasmeen K. Farouk. **Bioinformatics and Genomics Analytics:** Ahmed Elmahy, Ahmed El-Hosseiny, Ahmed Moustafa, Ashraf M. Muhammad, Asmaa Ali, Eman Adel, Enas Fouad, Khlood R. AbdElaal, Mahynour Albarbary, Sheri Saleeb, Tasnim A. Ghanim, Usama Bakry, Yasmeen Howeedy. **Clinical Genomics:** Alaa Hassan, Amira Kotb, Ayten Abdelaal, Eman Ramadan, Mohamed A. Elmonem, Neemat Kassem. **Information Technology:** Mahmoud El Ghoneimi, Mahmoud Fawzi, Mahmoud Samir.

**Sample Collection and Data Entry.** Aisha Mohamed Gamal-Eldin Abd El Mageed (Shefaa Al-Orman Hospital); Azza Saleh (Theodore Bilharz Institute; Shefaa Al-Orman Hospital); Engy A. Ashaat (NRC); Hadeer Elsaeed Abo-Elfarh (Mansoura University); Mohamed El-Gamal (Mansoura University); Naglaa Kholousy (NRC); Naglaa Zayed (Cairo University); Rania Zahwo (Alexandria University); Rehab Mohamed Ahmed (Shefaa Al-Orman Hospital); Shaimaa Ammar (Alexandria University); Sherif Abdel-Ghafar Fawzy (NRC); Shrouk Ahmed Muhamad (Mansoura University); Yasmine Mohamed Gaber (Cairo University); Zainab Wafik Zakaria Masoud (Cairo University).

## AI Disclosure

AI tools were used to assist with manuscript copyediting. The authors reviewed and edited all output and take full responsibility for the content of this publication.

## Author Contributions

**Khaled Amer:** Conceptualization, Data curation, Funding acquisition, Investigation, Methodology, Project administration, Resources, Supervision, Validation, Writing – original draft, Writing – review & editing. **Ahmed Moustafa:** Conceptualization, Data curation, Formal analysis, Investigation, Methodology, Software, Supervision, Validation, Visualization, Writing – original draft, Writing – review & editing. **Wael A. Hassan:** Data curation, Formal analysis, Investigation, Methodology, Project administration, Resources, Supervision, Writing – review & editing. **Eman Adel:** Data curation, Formal analysis, Investigation, Methodology, Software, Validation, Visualization, Writing – original draft, Writing – review & editing. **Khlood R. AbdElaal:** Data curation, Formal analysis, Investigation, Methodology, Software, Validation, Visualization, Writing – original draft, Writing – review & editing. **Tasnim A. Ghanim:** Data curation, Formal analysis, Investigation, Methodology, Software, Validation, Visualization, Writing – original draft, Writing – review & editing. **Ahmed Abd El-Raouf:** Formal analysis, Methodology, Writing – review & editing. **Ahmed El-Hosseiny:** Data curation, Formal analysis, Investigation, Methodology, Software, Validation, Visualization, Writing – review & editing. **Ahmed F. El-Sayed:** Investigation, Writing – review & editing. **Ahmed H. Badr:** Investigation, Methodology, Writing – review & editing. **Alaa Hassan:** Investigation, Writing – review & editing. **Amira Kotb:** Investigation, Writing – review & editing. **Amira Ragheb:** Investigation, Project administration, Writing – review & editing. **Ashraf M. Muhammad:** Data curation, Formal analysis, Investigation, Methodology, Validation, Visualization, Writing – review & editing. **Asmaa Ali:** Data curation, Formal analysis, Investigation, Methodology, Validation, Writing – review & editing. **Ayten Abdelaal:** Investigation, Writing – review & editing. **Eman Ramadan:** Investigation, Writing – review & editing. **Fadya M. El-Garhy:** Investigation, Methodology, Writing – review & editing. **Hasnaa El Shehaby:** Investigation, Methodology, Writing – review & editing. **Mahmoud A. Ali:** Investigation, Writing – review & editing. **Mahynour Albarbary:** Data curation, Formal analysis, Investigation, Methodology, Writing – review & editing. **Marwa A. Zahra:** Investigation, Methodology, Writing – review & editing. **May Amer:** Investigation, Methodology, Writing – review & editing. **Mohamed A. Elmonem:** Investigation, Writing – review & editing. **Nermeen T. Fahmy:** Investigation, Methodology, Writing – review & editing. **Omnia M. Abdel-Haseeb:** Investigation, Methodology, Writing – review & editing. **Tokka M. Hassan:** Investigation, Methodology, Writing – review & editing. **Yara A. Daoud:** Investigation, Methodology, Writing – review & editing. **Yasmeen Howeedy:** Formal analysis, Investigation, Methodology, Writing – review & editing. **Yasmeen K. Farouk:** Investigation, Methodology, Writing – review & editing. **Sameh Soror:** Conceptualization, Methodology, Writing – review & editing. **Gina El-Feky:** Funding acquisition, Project administration, Writing – review & editing. **Mahmoud Sakr:** Conceptualization, Funding acquisition, Resources, Writing – review & editing. **Neveen A. Soliman:** Conceptualization, Data curation, Investigation, Resources, Supervision, Writing – review & editing. **Yehia Z. Gad:** Conceptualization, Investigation, Methodology, Supervision, Writing – review & editing. **Khaled A. Abdel-Ghaffar:** Resources, Supervision, Writing – review & editing. **Egypt Genome Consortium:** Investigation, Resources.

