## Supplementary Materials for "EGP1K: Whole-Genome Sequencing of 1,024 Egyptians Characterizes Population Structure and Genetic Diversity"

### Supplementary Figures

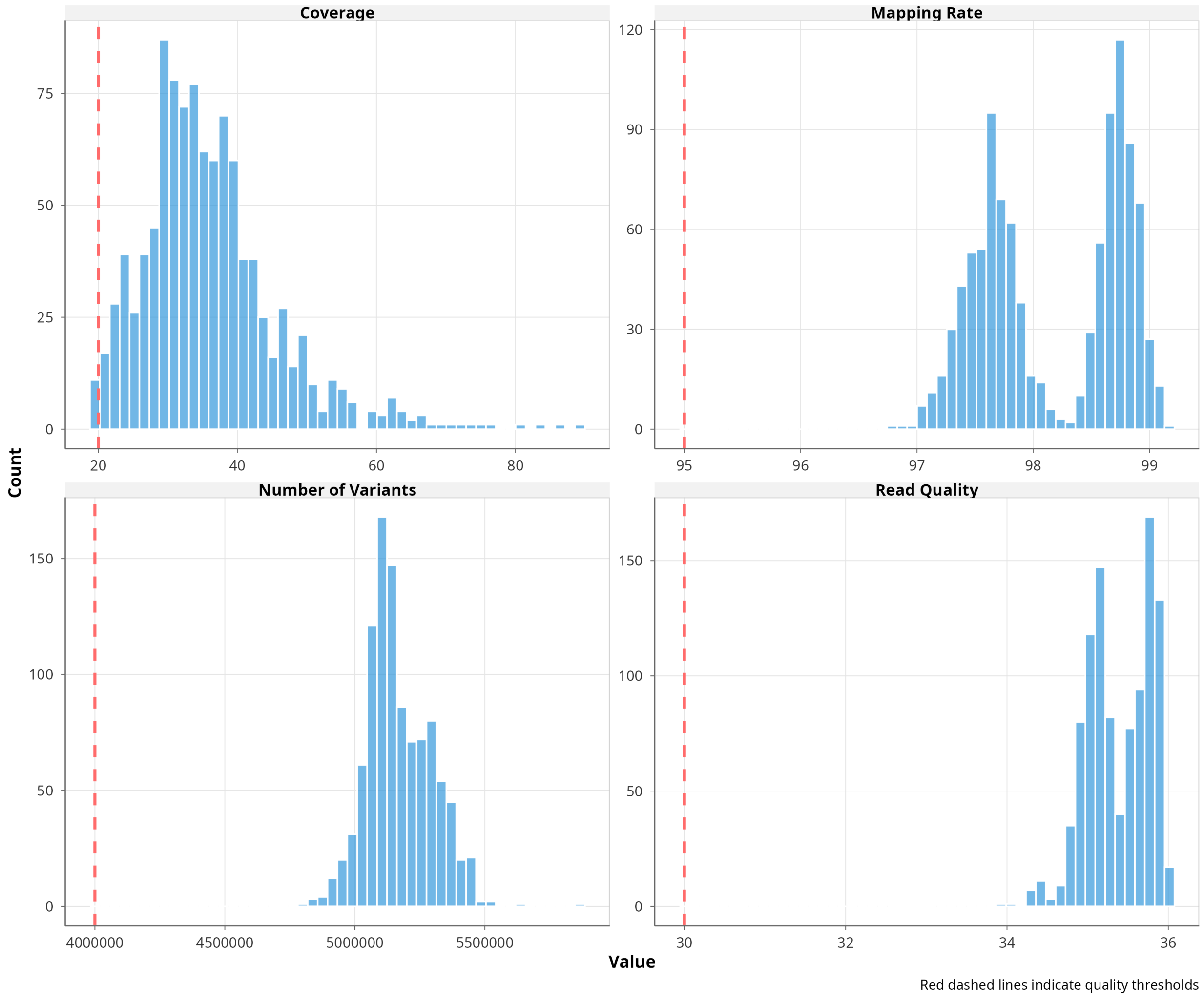


**Figure S1.** Distribution of sequencing quality metrics. Histograms showing the distribution of (A) sequencing coverage, (B) mapping rate, (C) variants per sample, and

(D) read quality (Phred score) across all 1,024 individuals.

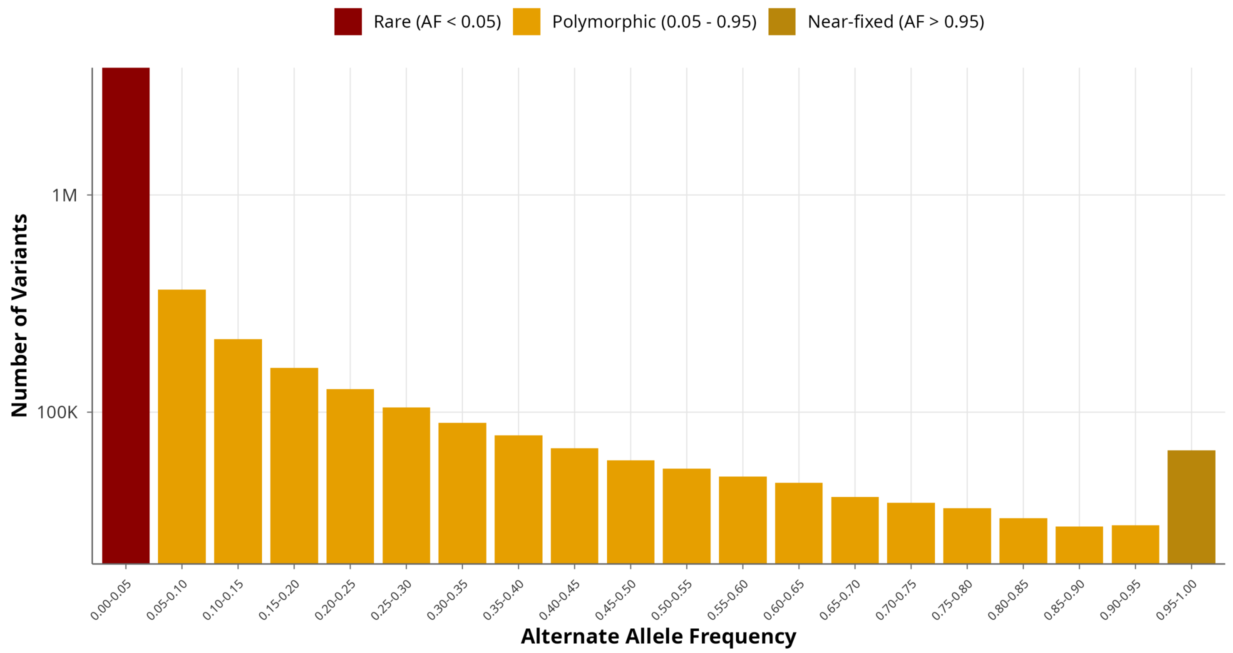


**Figure S2.** Site frequency spectrum of Egyptian autosomal variants. Distribution of alternate allele frequencies across 13.6 million autosomal variants identified in 1,024 Egyptian individuals. Bars are colored by frequency category: rare (AF below 0.05), polymorphic (AF 0.05 to 0.95), and near-fixed (AF above 0.95). The y-axis is shown on a logarithmic scale.

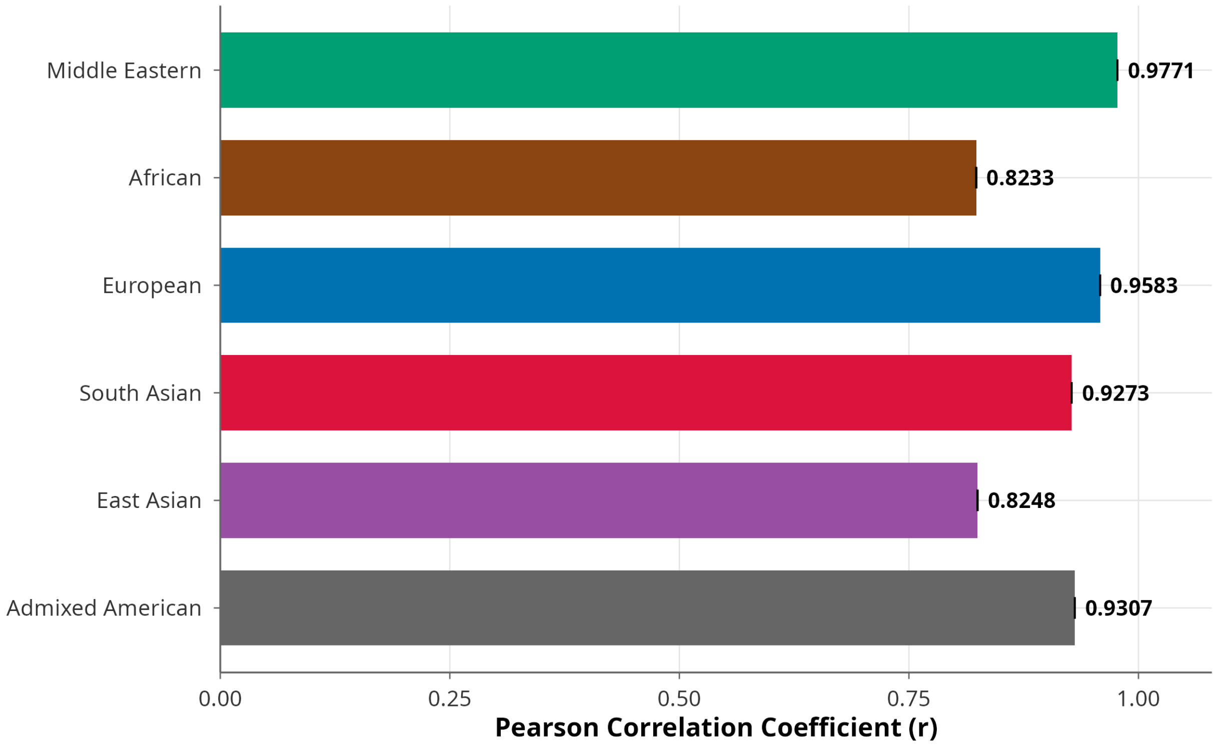


**Figure S3.** Allele frequency correlations between Egyptian and reference populations. Pearson correlation coefficients computed on a common intersection of 6,495,768 biallelic SNPs (MAF >= 0.01) with non-missing data across all seven populations. Confidence intervals are narrower than the bar widths due to the large number of variants.

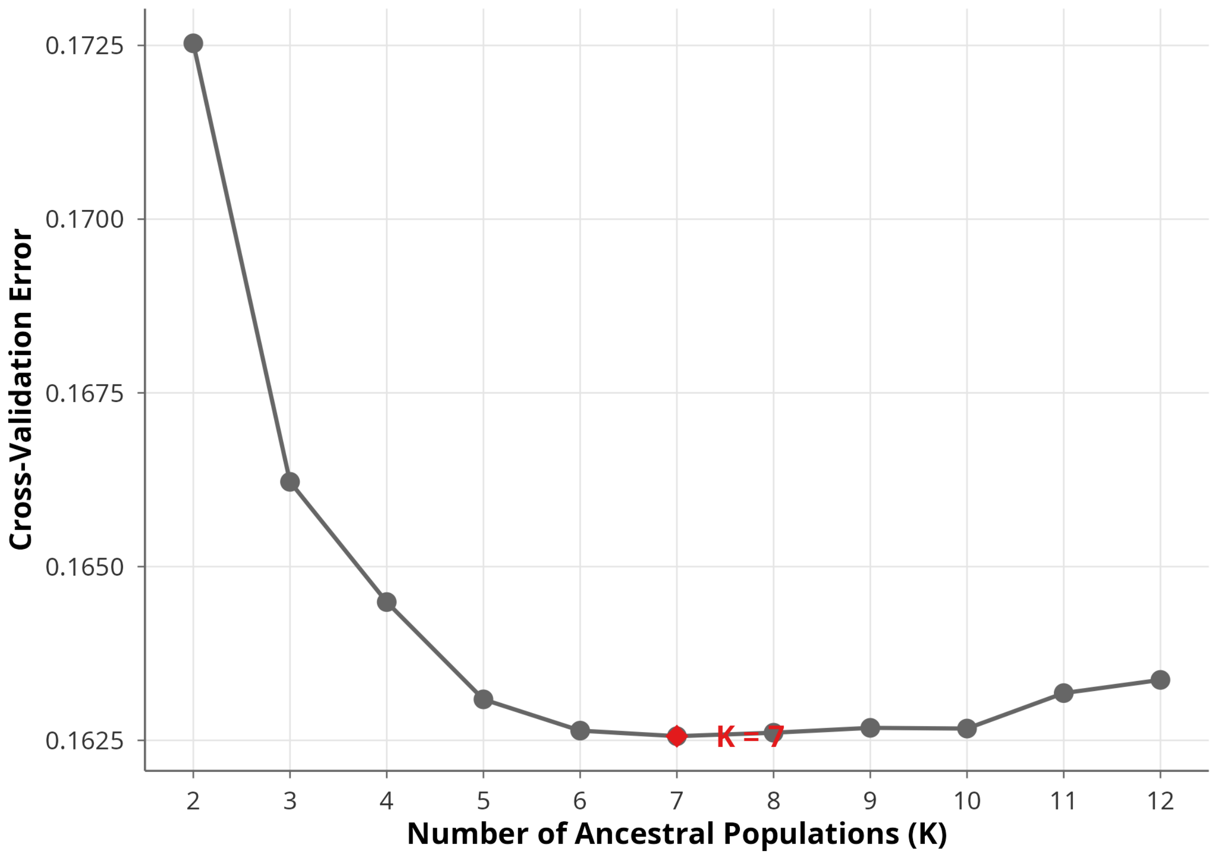


**Figure S4.** ADMIXTURE cross-validation error across K values. Cross-validation error for K = 2 through K = 12, computed from 5-fold cross-validation on genome-wide LD-pruned SNPs ($n$ = 29,138). The minimum at K = 7 (red diamond) was selected as the optimal number of ancestral clusters.

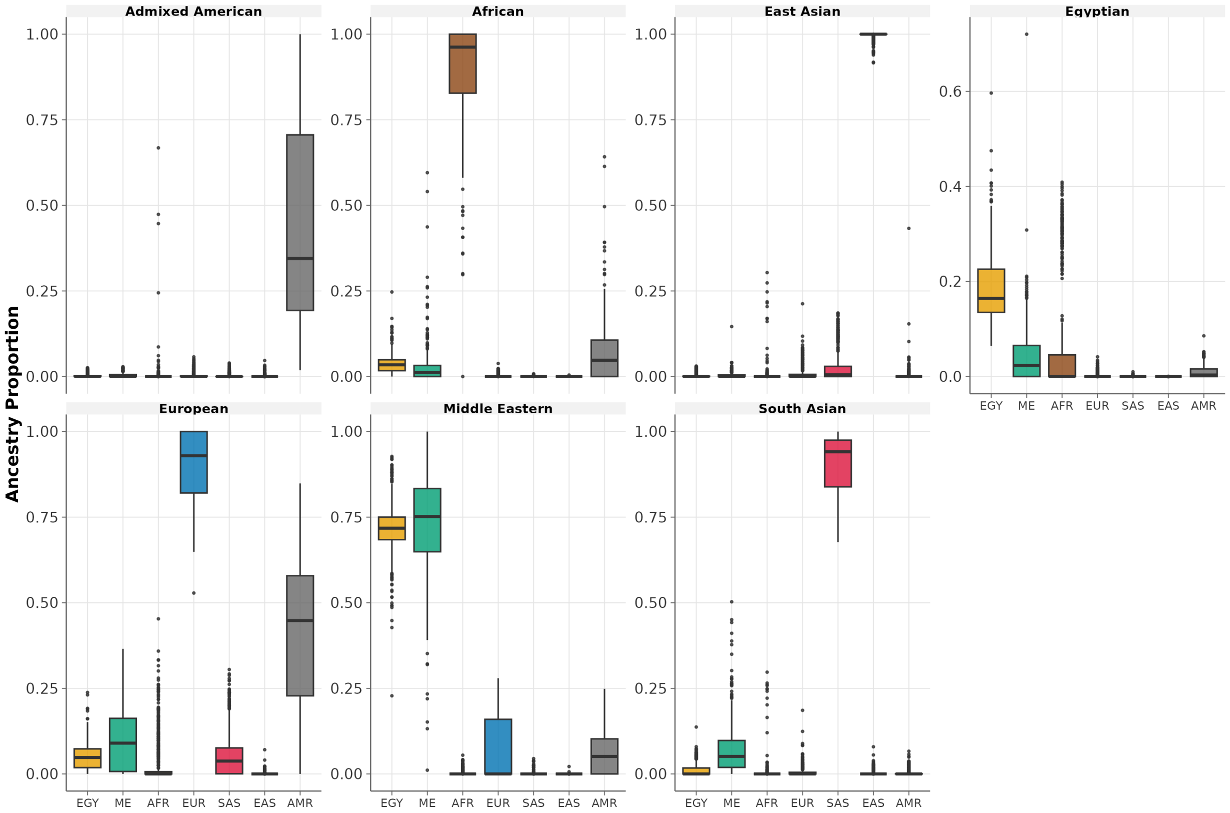


**Figure S5.** Distribution of ancestry proportions across populations at K = 7. Boxplots showing the distribution of each of seven ancestry components across Egyptian, Middle Eastern, African, European, South Asian, East Asian, and Admixed American populations.

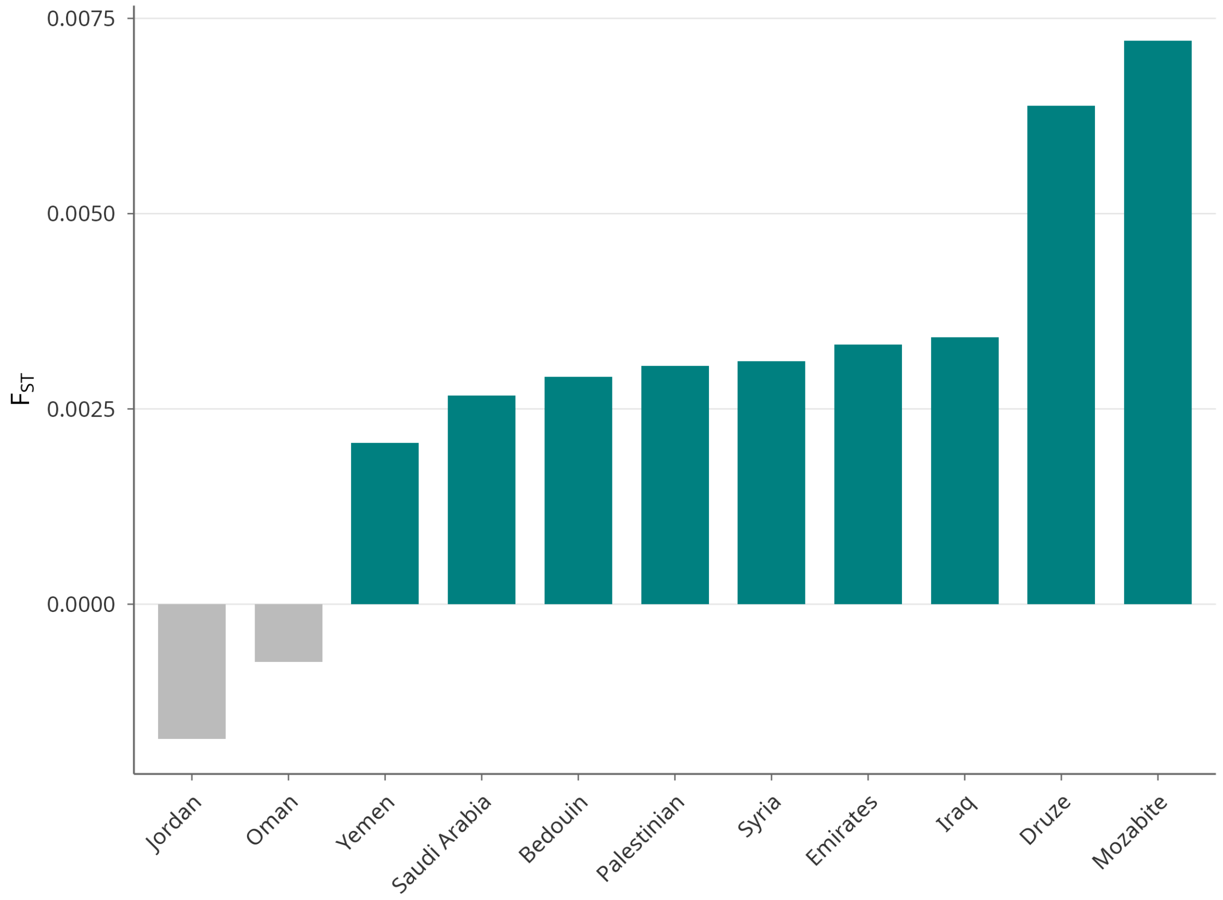


**Figure S6.** Pairwise Weir-Cockerham $F_{ST}$ between Egyptians and Middle Eastern subpopulations. Bar chart showing $F_{ST}$ values ordered by genetic distance. Yemeni (0.0021), Saudi (0.0027), and Bedouin (0.0029) show the lowest differentiation. Druze (0.0064) and Mozabite (0.0072) show the highest. Negative values for Jordan and Oman reflect sampling noise due to very small sample sizes ($n$ = 3 each) and should be interpreted as approximately zero differentiation.

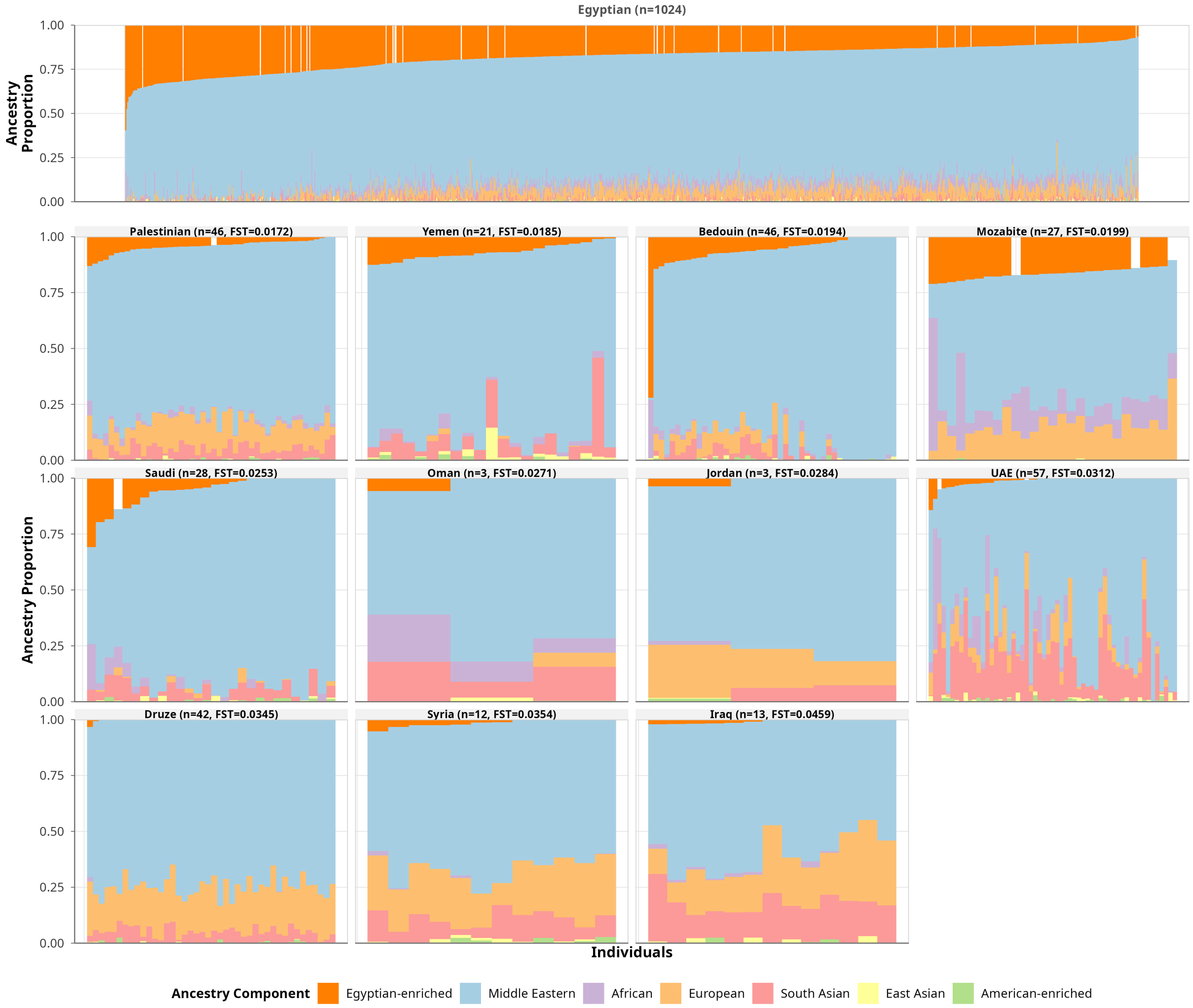


**Figure S7.** ADMIXTURE barplots for Egyptian and Middle Eastern subpopulations at K = 7. Panels are ordered by ADMIXTURE-based $F_{ST}$ distance from Egyptians, with the closest populations at the top and most differentiated at the bottom. Each vertical bar represents an individual; colors indicate ancestry proportions from seven clusters. Sample sizes and ADMIXTURE-based $F_{ST}$ values are shown for each panel. Note that sample sizes may differ from those in Figure S6 (Weir-Cockerham $F_{ST}$) due to variant-level filtering differences between the two analyses; in particular, Jordan is represented by $n$ = 1 individual in the ADMIXTURE analysis compared to $n$ = 3 in the Weir-Cockerham $F_{ST}$ analysis.

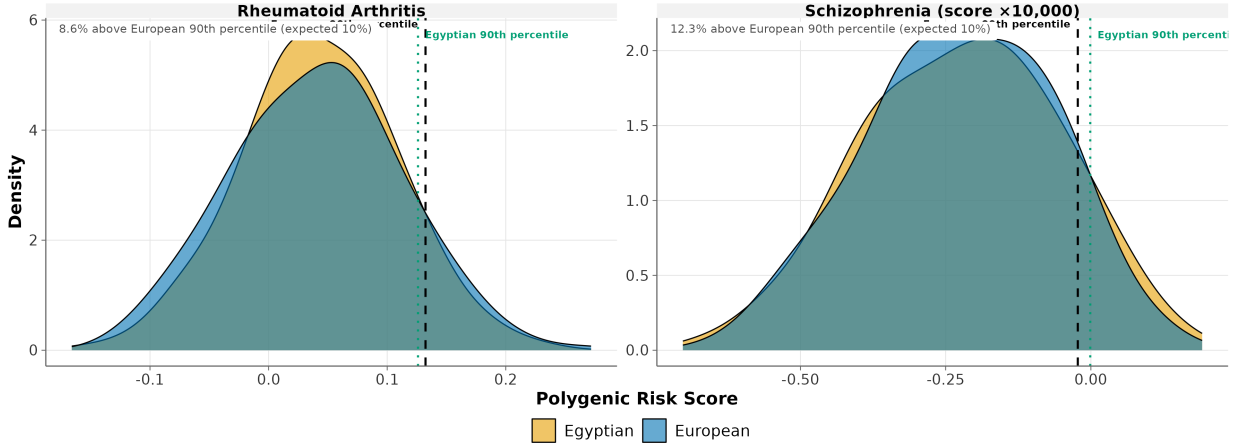


**Figure S8.** Examples of traits with limited cross-population PRS distributional shift. Density distributions of polygenic risk scores for rheumatoid arthritis and schizophrenia in Egyptian and European populations. Vertical lines indicate the European and Egyptian 90th percentile thresholds. In contrast to the cardiometabolic traits shown in Figure 7, these examples demonstrate close alignment of PRS distributions and high-risk thresholds across populations, with proportions of Egyptians exceeding the European threshold (8.6% and 12.3%) approximating the expected 10%.

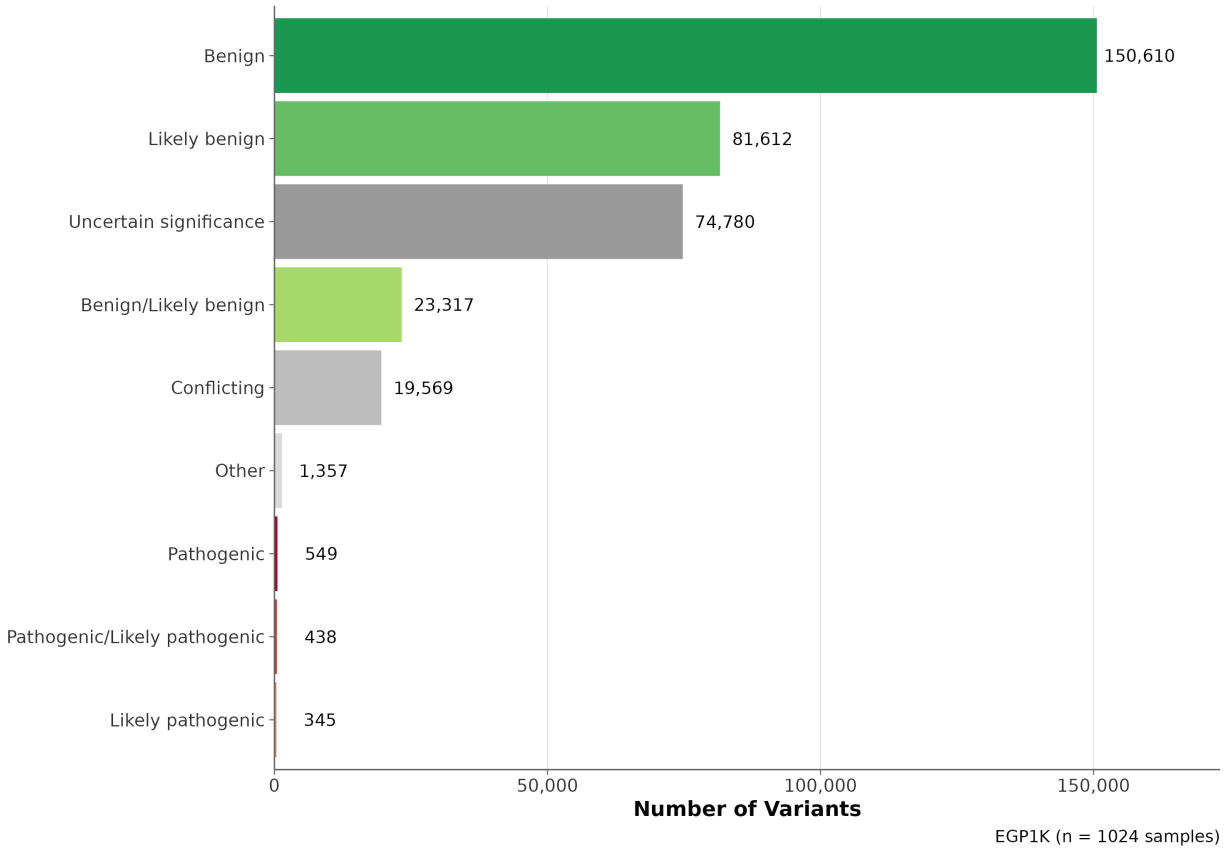


**Figure S9.** Distribution of ClinVar clinical significance classifications. Horizontal bar chart showing the distribution of 352,577 EGP1K variants at positions annotated in ClinVar, categorized by clinical significance.

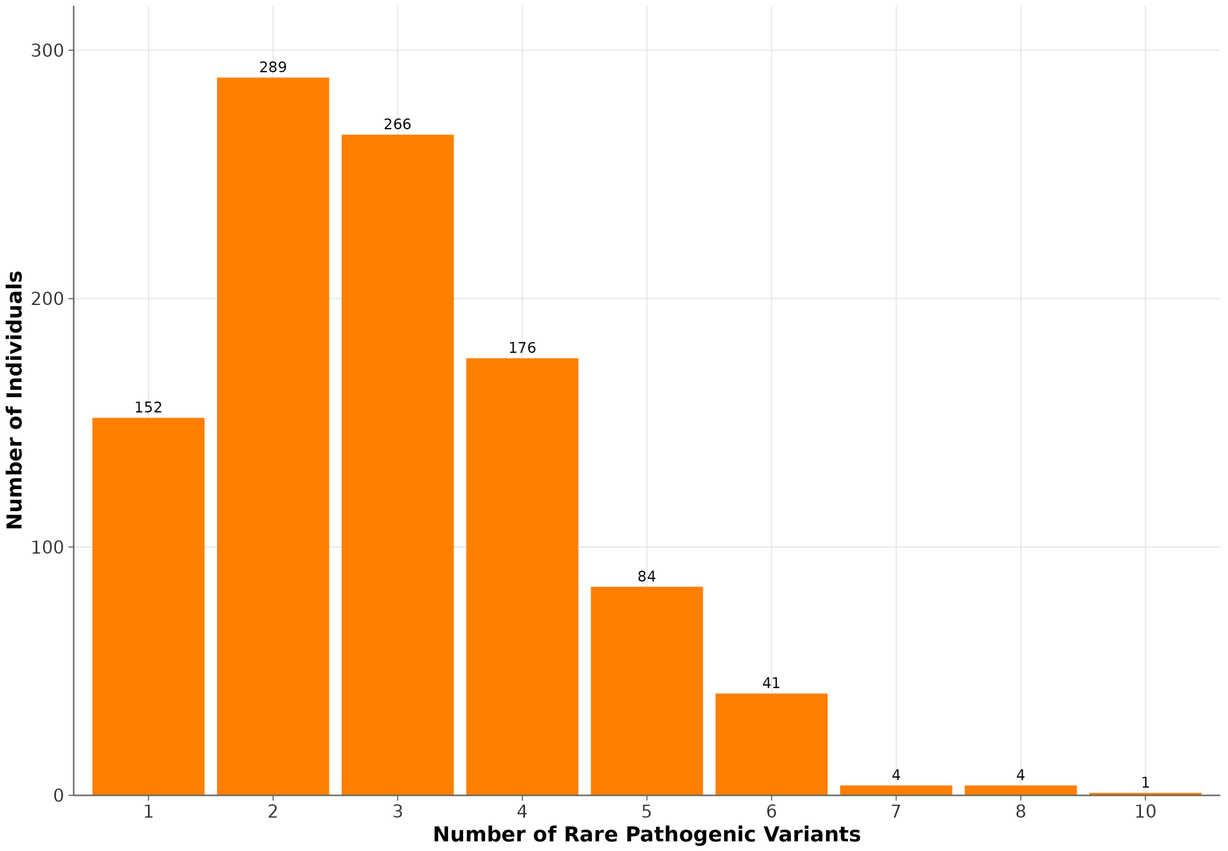


**Figure S10.** Distribution of rare pathogenic variant burden per individual. Number of rare pathogenic or likely pathogenic variants carried by each of 1,024 Egyptian participants.

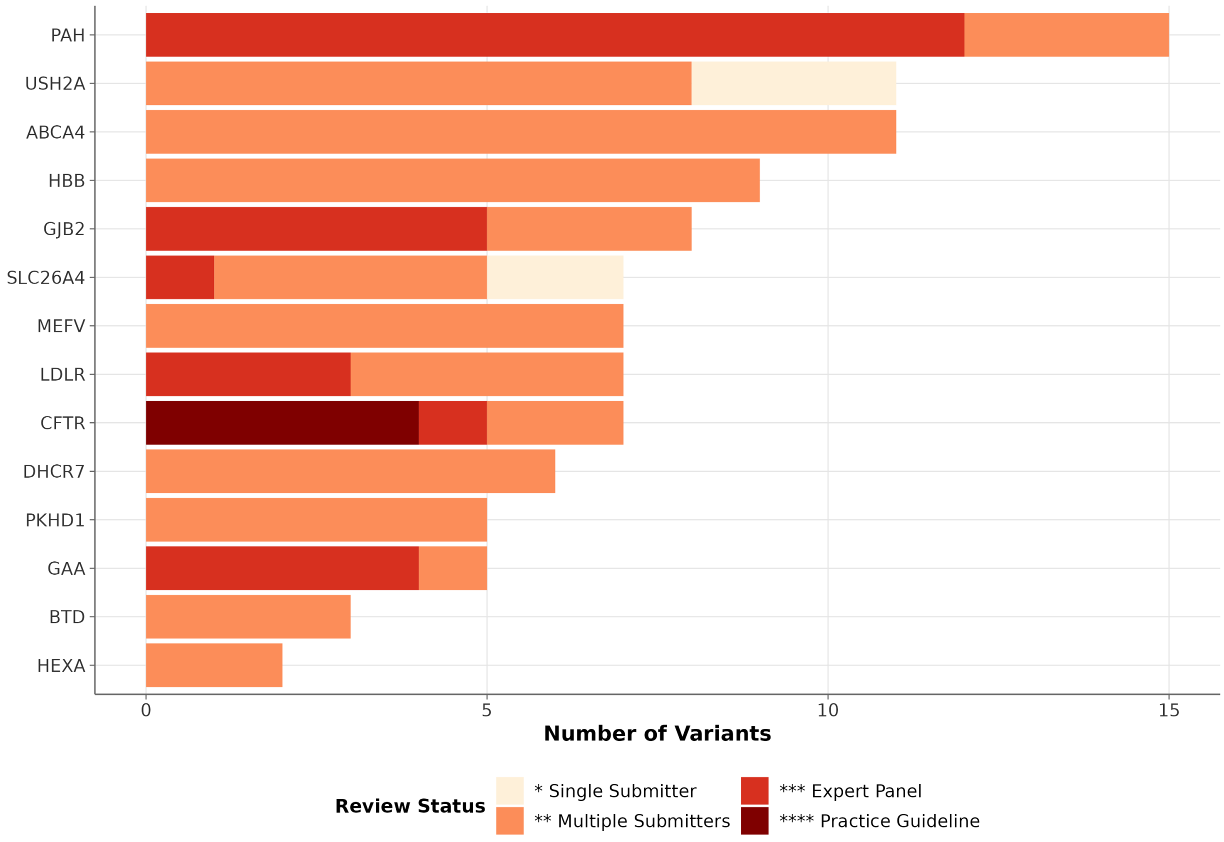


**Figure S11.** ClinVar evidence quality for pathogenic variants by gene. Stacked bar chart showing the distribution of ClinVar review status (star ratings) for pathogenic and likely pathogenic variants across 14 disease genes.

### Supplementary Tables

**Table S1.** Sequencing quality metrics for 1,024 Egyptian individuals. Summary statistics: mean coverage 35.9x (SD 9.7x; range 19.5-89.3x), mean mapping rate 98.17% (SD 0.6%), mean read quality Q35.4. Full per-sample data provided in the accompanying TSV file (Table_S1_Sequencing_Quality_Metrics.tsv).

**Table S2.** Pathogenic MEFV variants identified in the Egyptian cohort.

| Variant | Heterozygous | Homozygous | Total Individuals | Het Carrier Freq (%) |
|---|---|---|---|---|
| chr16:3243310 A>G (M694V) | 48 | 2 | 50 | 4.69 |
| chr16:3243405 C>T (V726A) | 29 | 0 | 29 | 2.83 |
| chr16:3243447 C>T (M680I) | 9 | 0 | 9 | 0.88 |
| chr16:3243407 T>C | 4 | 0 | 4 | 0.39 |
| chr16:3243205 C>T | 3 | 0 | 3 | 0.29 |
| chr16:3243408 CATT>C | 1 | 0 | 1 | 0.10 |
| chr16:3243529 C>T | 1 | 0 | 1 | 0.10 |

All variants classified as Pathogenic or Likely Pathogenic in ClinVar (version 2024-11). "Total Individuals" counts all individuals carrying the variant (heterozygous and homozygous). "Het Carrier Freq" is calculated as heterozygous carriers divided by 1,024 unrelated individuals. Individuals who are compound heterozygous for two MEFV variants (Table S3) are counted once per variant in the heterozygous column. Allele counts in the full data file may exceed the expected value of n_het + 2 * n_hom for some variants due to multi-allelic site representation in the merged VCF.

**Table S3.** Homozygous and compound heterozygous carriers of pathogenic variants.

| Sample | Gene | Disease | Category | Variants |
|---|---|---|---|---|
| EGP_0358 | MEFV | Familial Mediterranean Fever | Homozygous | M694V (1/1) |
| EGP_0792 | MEFV | Familial Mediterranean Fever | Homozygous | M694V (1/1) |
| EGP_0979 | MEFV | Familial Mediterranean Fever | Compound heterozygous | M694V + M680I |
| EGP_0257 | MEFV | Familial Mediterranean Fever | Compound heterozygous | M694V + V726A |
| EGP_0628 | CFTR | Cystic Fibrosis | Homozygous | chr7:117548630 T>G (1/1) |
| EGP_0031 | GAA | Pompe Disease | Compound heterozygous | chr17:80104542 T>G + chr17:80108547 C>G |
| EGP_0843 | GAA | Pompe Disease | Compound heterozygous | chr17:80104542 T>G + chr17:80112679 G>A |

**Table S4.** Hardy-Weinberg equilibrium test results for pathogenic variants. All tested variants were in Hardy-Weinberg equilibrium after FDR correction (all adjusted $p > 0.05$). Testing was performed on variants with sufficient allele counts for exact tests; genes with very low carrier counts (fewer than 5 carriers) were not assessed. Full test statistics provided in the accompanying TSV file (Table_S4_HWE_Test_Results.tsv).

**Table S5.** Published carrier frequency reference values used for population comparisons in Figure 9. Full reference table with source citations provided in the accompanying TSV file (Table_S5_Reference_Carrier_Frequencies.tsv).

### Supplementary Note 1: Detailed Variant Discovery Statistics

Sequencing achieved a mean coverage of 35.9x (SD = 9.7x; range: 19.5-89.3x), a mean mapping rate of 98.17% (SD = 0.6%), and a mean read quality of Q35.4 (Table S1; Figure S1). On average, each participant carried 3,844,588 variant sites (heterozygous or homozygous alternate). The mean total variant count per sample, including homozygous reference calls at variable positions, was 5.17 million. When restricted to autosomal biallelic variants, 13.6 million sites were polymorphic, of which 10,765,485 (79.0%) had a minor allele frequency below 1%.

Most variants were located in intergenic (41.01%) and intronic (49.10%) regions, whereas only 1.75% occurred within exonic regions. Among exonic variants, the largest categories were nonsynonymous (54.5%), synonymous (40.3%), and frameshift (1.2%), with the remainder comprising stopgain, stoploss, and non-frameshift indels. In total, 274,607 singleton variants (alternate allele count = 1 across all 1,024 Egyptian individuals) were identified, corresponding to an average of 268 singleton sites per individual.

Starting from 82,804,613 raw variants, staged quality control through GATK VQSR (reducing to 64,835,610 variants), restriction to GIAB HG001 high-confidence regions (52,100,628 variants), and functional annotation produced the final set of 51,342,784 variants.

### Supplementary Note 2: Detailed HLA Class I Allele Frequencies

At the HLA-A locus, A*02:01 was the most prevalent allele at 7.10%, followed by A*01:01 (5.80%), A*30 (3.60%), A*24 (2.90%), and A*03 (2.50%). At the HLA-B locus, B*41:01 and B*35:01 showed nearly equal frequencies at 3.80% and 3.70%, respectively, followed by B*52 (2.20%), B*14 (2.10%), and B*15 (2.10%). At the HLA-C locus, C*07:01 was the most common allele at 6.90%, followed by C*12:01 (5.10%), C*04 (4.90%), C*17 (3.80%), and C*06 (3.80%). Alleles are reported at the highest resolution achieved by the DRAGEN HLA Caller; some alleles could only be resolved to one-field (two-digit) level.
